# Systematic hyper-variation and evolution at a lipopolysaccharide locus in the population of *Xanthomonas* species that infect rice and sugarcane

**DOI:** 10.1101/2022.03.02.482478

**Authors:** Anu Singh, Kanika Bansal, Sanjeet Kumar, Prabhu B. Patil

## Abstract

Advent of high throughput sequencing and population genomics is enabling researchers to investigate selection pressure at hyper-variable genomic loci encoding pathogen-associated molecular patter (PAMP) molecules like lipopolysaccharide (LPS) in an unprecedented manner. *Xanthomonas* is a model group of phytopathogenic bacteria that infects host in tissue-specific manner. Our in-depth investigation revealed that the successful emergence of lineages infecting major cereals and grasses like rice, sugarcane, and wheat was mediated by acquisition and later replacement of an ancestral type (BXO8) of LPS cassette by distinct one. In the population of the rice xylem pathogen, *X. oryzae* pv. *oryzae* (Xoo), the BXO8 is replaced by a distinct BXO1 type of cassette. Alternatively, in diverse *Xanthomonas* species that infect sugarcane, the BXO8 ancestral cassette has been replaced by yet another kind of Xvv type of LPS cassette, suggesting convergent evolution at an LPS locus mediated by horizontal gene transfer (HGT) events. Aside from xylem, two closely related lineages of *X. oryzae* that infect parenchyma tissue of rice and *Leersia hexandra* grass have acquired an LPS cassette from *Xanthomonas* pathogens that infect citrus, walnut, and strawberry parenchyma, indicating yet another instance of parallel evolution facilitated by HGT. Our targeted and mega-population-based genome dynamic studies revealed potential role of acquisition of specific types of LPS cassettes in the emergence and evolution of tissue specificity in *Xanthomonas*. Additional cellular, molecular, genetic, and plant studies will help us figure out how a distinct type of LPS help *Xanthomonas* pathovars and lineages adapt to parenchyma and xylem tissues.

## Introduction

Phytopathogen genomics and systemic-functional studies have provided a remarkable understanding of host-microbiome interactions, virulence, and host adaptation pathways, giving significant insights into microbial ecology and epidemiology (An et al., 2020). LPS is a major component of the outer membrane of phytopathogens that is well known to act as a pathogen-associated molecular pattern (PAMP), virulence determinant, and an elicitor of defence responses (Clifford, Rapicavoli, & Roper, 2013). It accounts for nearly 75% of the bacterial cell surface and most likely is a pivotal contributor to the adhesion of the bacterial cell to the host cells (Alexander & Rietschel, 2001; Goldberg & Pler, 1996; Walker, Redman, & Elimelech, 2004). Interestingly, it has a significant role as a stimulator of the immune system in both humans and plants. LPS possess long O-antigen that restrict the initial plant recognition, thereby enabling elicitation of innate immunity and helping to evade successfully into the host (Ranf et al., 2015; Rapicavoli et al., 2018). Whereas the recent plant immunology study also demonstrated that acetylated LPS in *Arabidopsis thaliana* protects the LPS from immune recognition (Vanacore et al., 2022), and it is found to activate biphasic production of reactive oxygen species (ROS) (Shang-Guan et al., 2018). In addition, LPS has been discovered to be a fundamental factor that increases bacterial viability and aids in inter-organismic interaction as well. It increases bacterial virulence during host infection and distracts bacteria from host immune responses (Kutschera & Ranf, 2019). It has recently been shown to induce a defence-response in rice cells that leads to programmed cell death (Desaki et al., 2006). Surprisingly, LPS in human and pathogenic bacteria varies greatly across species and strains. With the availability of a large number of genome sequences, there is an exciting opportunity to understand the evolutionary pattern at the population level and selection pressure at LPS loci in pathogenic bacteria.

*Xanthomonas* is a gram-negative, highly evolved, and extremely successful group of phytopathogens that infects a wide range of dicot and monocot plants (NIÑO◻LIU, Ronald, & Bogdanove, 2006). *Xanthomonas* pathovars are also used as model pathogens to study the evolution of host specificity and adaptation in pathogen bacteria. These species encompasses a diverse range of plant pathogens that use various pathogenicity modes in order to survive and thrive in the host (Timilsina et al., 2020). *Xanthomonas* species are classified into two clades: the minor clade I and the major clade II (Bansal, Kumar, Kaur, Singh, & Patil, 2021). Clade I represents early branching and comprises pathogens with reduced genome such as *X. albilineans* (Xalb) (Mensi, Vernerey, Gargani, Nicole, & Rott, 2014) as well as non-pathogens like *X. sontii* (Bansal, Kaur, et al., 2021) whereas the clade II comprises pathogens such as *X. oryzae* and *X. citri* (Bansal, Midha, Kumar, & Patil, 2017; Midha et al., 2017). While most clade II species infect dicot parenchyma, such as *X. citri* and *X. arboricola*, a few exceptions to this clade are monocot infecting *X. oryzae*, *X. axonopodis*, and *X. vasicola*, which are well-known pathogens of cereals and grasses. Clade I, on the other hand, is made up of species that primarily infect monocots, such as *X. sacchari* and *X. albilineans*. While Clade II is composed of *X. oryzae* pv. *oryzae* or Xoo the causal agent of bacterial leaf blight, that infects the xylem of rice plants, a staple crop for more than half the world population. There are two more pathovars of *X. oryzae* that infect parenchyma tissue. One is *X. oryzae* pv. *oryzicola* (Xoc) that causes bacterial leaf streak in rice, and the other is *X. oryzae* pv. *leersia* (Xol) that infects *Leersia hexandra* grass. One of its two major relatives are *X. axonopodis* (Xaxn), which comprises of two groups: *X. axonopodis* that infects grasses (Constantin et al., 2016) and the other is *X. axonopodis* pv. *vasculorum* that infects sugarcane. Along with this, clade II also consists of *X.* vasicola pv. *vasculorum* (Xvv) which infects sugarcane, and causes gumming and leaf chlorotic streak disease and *X. citri pv. citri* (Xcc), infecting primarily fruit crops and citrus (Patané et al., 2019). Clade I consist of *X. albilineans* (Xalb) and *X. sacchari* (Xsac) which are two closely related species infecting sugarcane. Xsac causes leaf chlorotic streak disease of sugarcane (Sun et al., 2017) and Xalb leads to leaf scald, a vascular disease of sugarcane (Muimba-Kankolongo, 2018), respectively. Previously, it was believed that the pathogen of Xalb was restricted to the xylem of sugarcane. However, it was detected in the parenchyma and bulliform cells of infected leaves. It is now known that the pathogen initially exists in the xylem and infects the parenchyma after creating openings by degrading the cell wall and middle lamellae. So, it is interesting to note that Xalb is a sugarcane pathogenic vascular bacterium that invades and multiplies in otherwise healthy non-vascular plant tissues (Mensi et al., 2014).

LPS is an essential factor in *Xanthomonas* pathogenicity and virulence, as it aids in evading host immune responses (Babu et al., 2014; Lang et al., 2019). An LPS locus between housekeeping genes *metB* and *etfA* in the genus *Xanthomonas* is hypervariable within species/strains (Patil & Sonti, 2004). Mutations at this locus have been linked to a loss of virulence in both Xoo and Xoc (Dharmapuri, Yashitola, Vishnupriya, & Sonti, 2001)(Wang, Vinogradov, & Bogdanove, 2013). It has been reported that the Xvv pathogen can infect sugarcane via LPS, which is similar to *X. axonopodis* pv. *citri* (Wasukira et al., 2014). However, studies on the LPS locus at the population level within and between pathovars are lacking. Earlier studies revealed three types of LPS gene clusters or cassettes (BXO1, BXO8, and BLS256 type) in *X. oryzae* (Patil & Sonti, 2004). Even within lineages of Xoo, there are two types of LPS cassettes which only infect rice xylem, suggesting LPS as a potential determinant of tissue-specificity. However, these and other studies on variation on LPS locus were based on a limited number of genomes, and with the explosion in the number of sequenced genomes of members of the genus *Xanthomonas*, there is a scope and need to carry out deeper and population-based comparative studies.

Currently, the NCBI genome portal contains 406 high-quality genomes from *X. oryzae*, submitted by various research groups around the world (Zheng et al., 2020). Apart from *X. oryzae*, the genomic resources of its relatives from other species are also abundant. As of December 2021, there were 151genomes of Xcc, 114 genomes of Xvv, 17 genomes of Xalb along with five genomes of Xaxn and four genomes of Xsac available on GTDB (https://gtdb.ecogenomic.org/) and NCBI portal. Hence, this provides an unprecedented opportunity to investigate the type of LPS cassettes found in *Xanthomonas* pathovars and their association with tissue specificities. There is also potential to investigate the association with host specificity by robustly comparing the variation with other members of the *Xanthomonas* genus with similar lifestyles using mega-population-based targeted comparative genomic studies. In this present study, we aim to conduct targeted and systematic comparative and evolutionary genomics at the population level in order to understand large scale variation at LPS loci.

## Results

### Evolutionary tracing reveals the presence of the BXO8 type of LPS cassette in diverse *Xanthomonas* species that infect the xylem of rice, carpet grass, and sugarcane

To understand the diversity, we screened the LPS locus for the type of LPS cassette variation available in the NCBI. The analysis revealed the presence of a 21 KB cassette with 57.1% GC content and encoding 17 ORFs, which we refer to as the BXO8 type LPS or grass type of LPS cassette. One part of the cassette harbours genes involved in LPS biosynthesis, and the remaining cassette harbours genes involved in virulence and accessory genes. In the BXO8 type LPS cassette, there is a set of nine genes at the *etfA* end involved in NAD/FAD dependent-oxidoreductase, prenyltransferase, methyltransferase, and GtrA family proteins involved in the synthesis of cell surface polysaccharides. The initial ORFs from the *metB* side of BXO8 were annotated as *wzm*, *wzt* genes, *glycosyltransferase*, and some hypothetical proteins.

Phylogenetic analysis also revealed that this intact BXO8 type LPS is present in strains belonging to four *Xanthomonas* species, i.e., Xoo, Xaxn, Xvv, and Xsac that infect the xylem of rice, grasses (*Axonopus scoparius* and *Axonopus micay*), and sugarcane, respectively (Figure 1A, Figure 1B). While it is interesting to note that Xoo, Xaxn, and Xvv belong to clade-II, Xsac belongs to clade-I. Population level genomic studies revealed that 19/406 strains of Xoo (figure 2A), 5/5 strains of Xaxn, 19/113 strains of Xvv (figure 3A), and 1/4 strains of Xsac (figure 4A) have BXO8 type of LPS cassette. Homology blast results for all the genomes is provided in supplementary table 1. We compared the cassette to interspecies variation and found minor differences at some IS elements and hypothetical genes (figure 1B). Minor differences between *X. axonopodis* pv. *vasculorum* NCPPB 796 (T) and BXO8 type LPS include the presence of one more phytanoyl-coA dioxygenase family protein, the presence of IS5 transposons, and the presence of two new hypothetical genes whose functions are unknown. In a similar way, in the case of the Xvv NCPPB 795 strain and *X. sacchari* F10, there is more than 90% query coverage and 80% above percent identity found with the BXO8 type of LPS cassette. The arrangement of genes from the both *etfA* and *metB* sides in all the genomes is similar with BXO8 (figure 1B).

**Figure 1:**
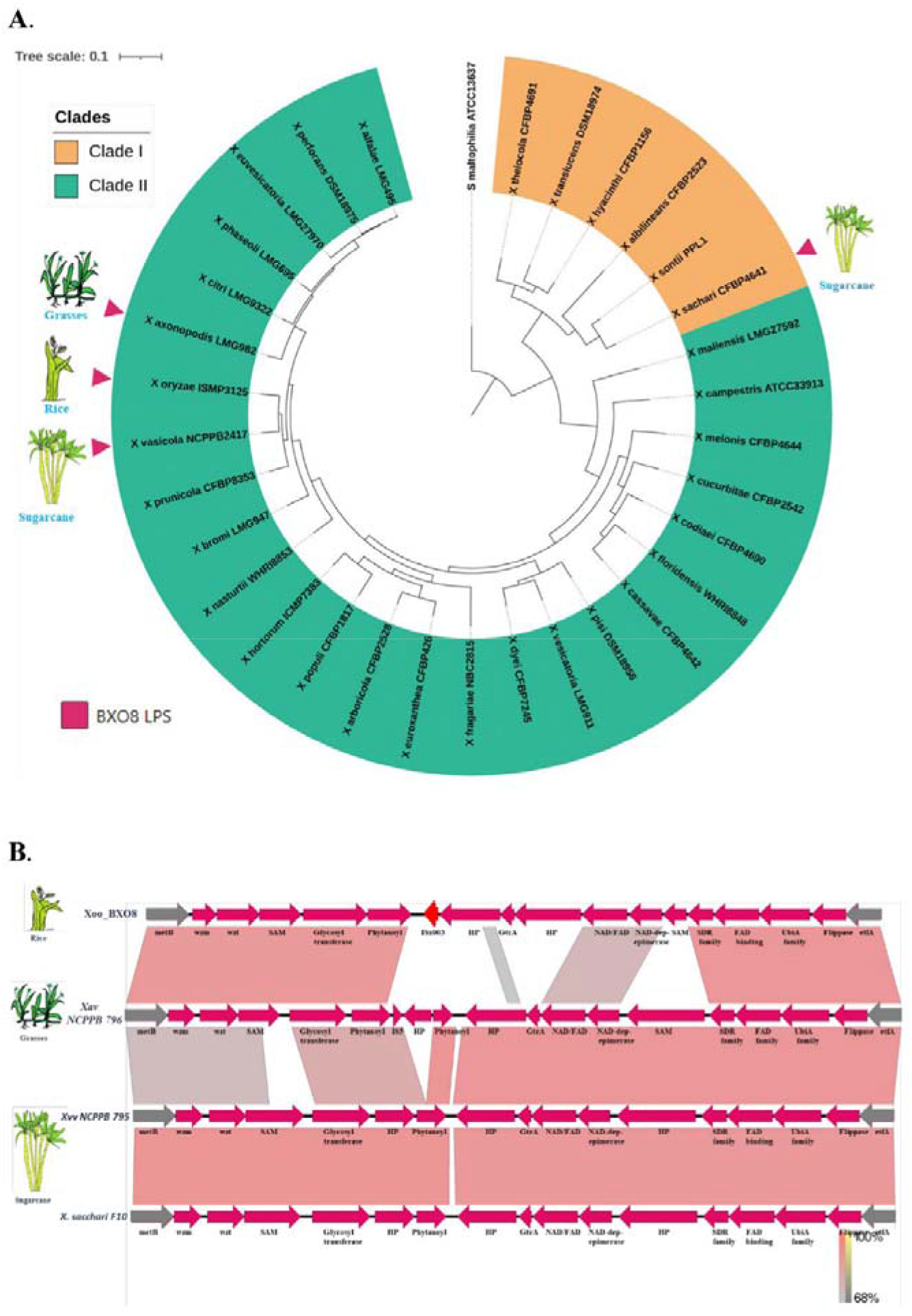
Phylogenetic tree of *Xanthomonas* type strains. (**A**) Fast-tree was used to reconstruct the core-gene tree, which was then visualised using iTOL software. Clade I depicted in yellow, while Clade II is depicted in green. The presence of BXO8 type LPS is indicated by a pink coloured arrow. (**B**). Easyfig and the BLASTn algorithm were used to compare the LPS synteny of Xoo-BXO8, Xac NCPPB 796, Xvv NCPPB 795, and X. sacchari F10. The gene locations are represented by arrows, and the degree of homology between pairs of genes in two LPS cassettes is represented by shaded lines. The *etfA* and *metB* genes are represented by grey arrows, while IS elements are shown in red.

**Figure 2:**
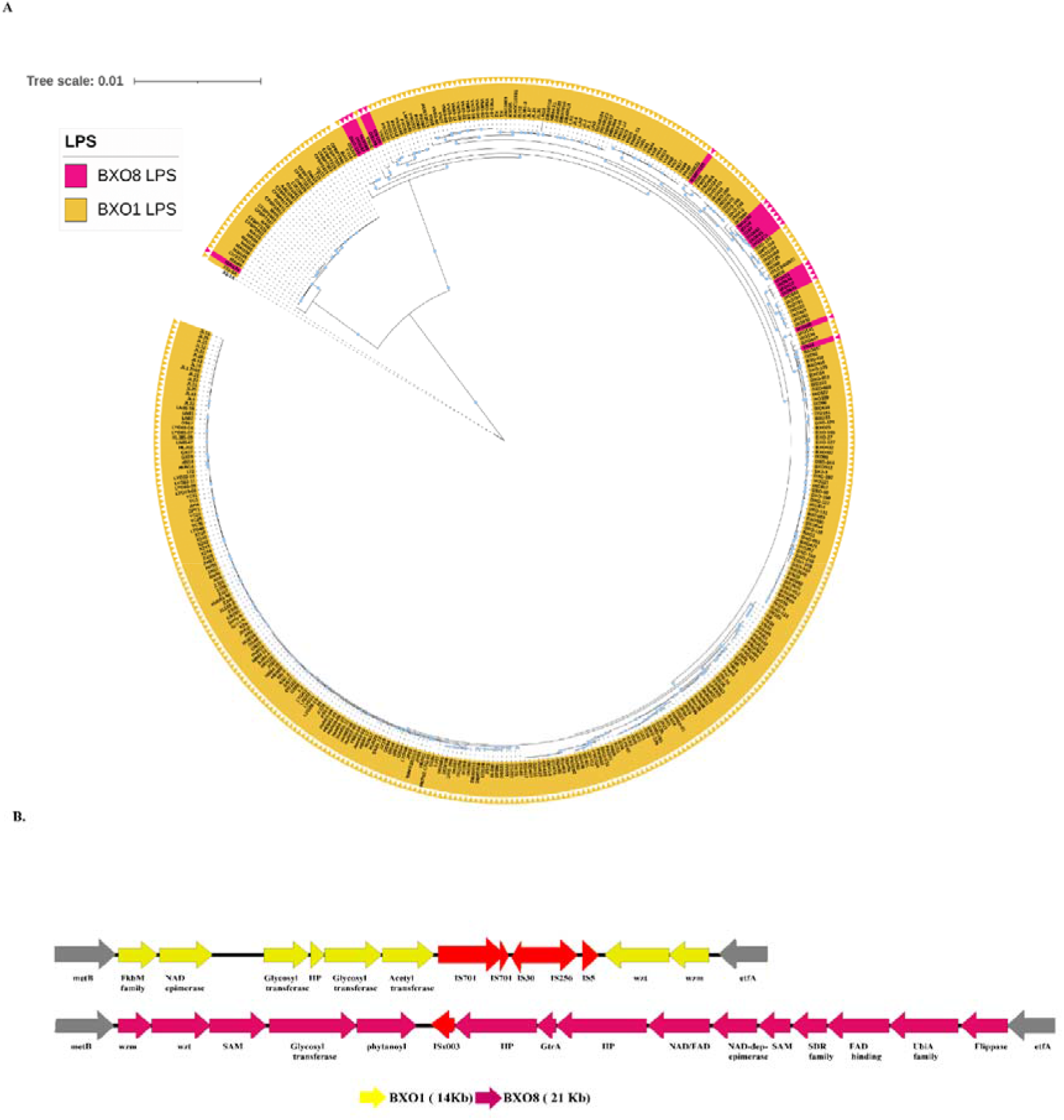
Distribution of LPS in the Xoo population. (**A**) PhyML was used to reconstruct the core-gene tree, which was then visualised using iTOL software. The yellow colour represents BXO1 type LPS, while the pink colour represents BXO8 type LPS found in each strain. The number of substitutions per site is indicated by the scale bar. (**B**). Genetic comparison of LPS cassettes of the BXO1 and BXO8 types. The location and direction of genes are represented by arrows, and homology between two genes is represented by similar colours. The *etfA* and *metB* genes are represented by grey arrows, while IS elements are shown in red.

**Figure 3:**
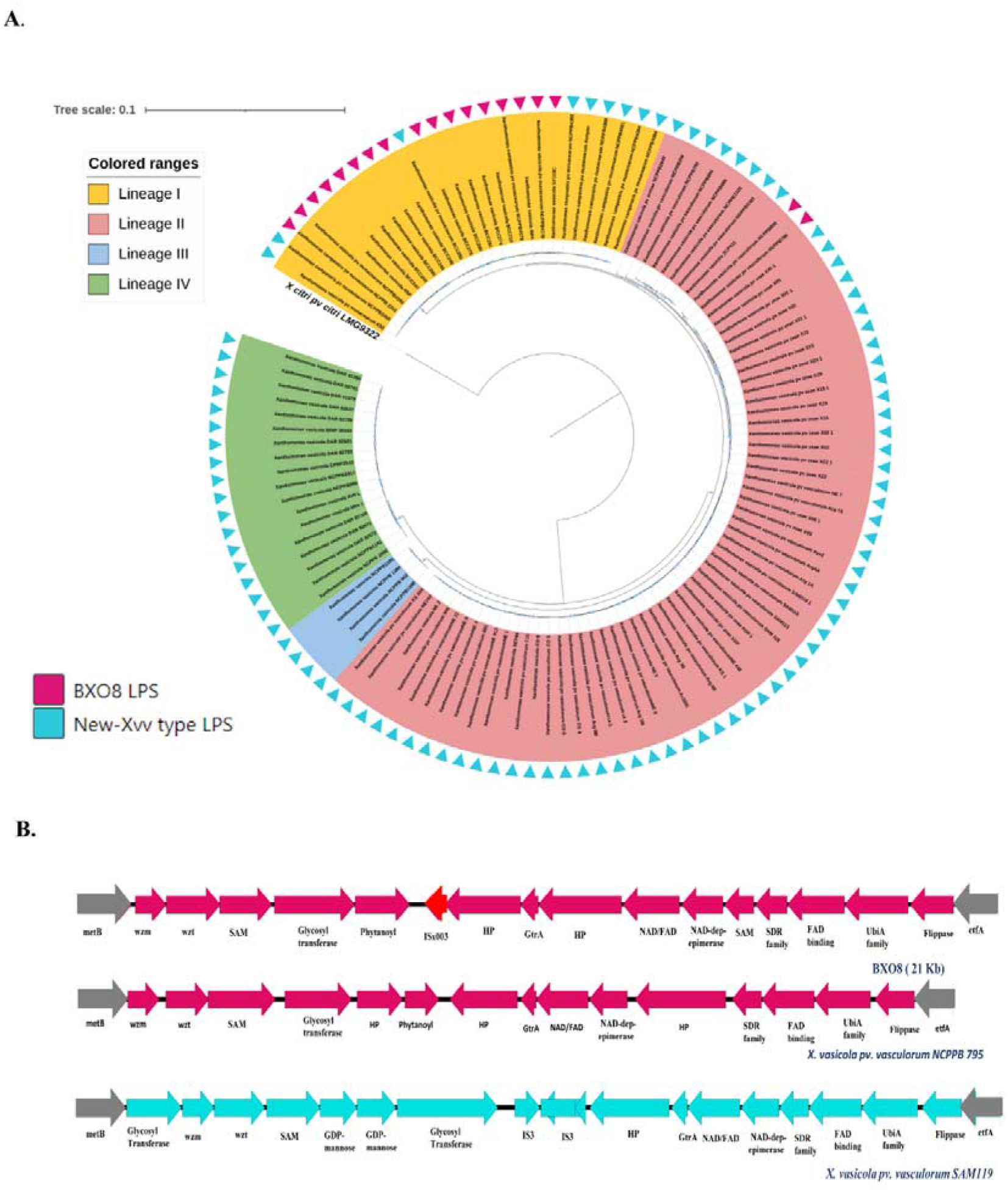
Population phylogenomics of *X. vasicola*. (**A**) Fast-tree was used to reconstruct the core-gene tree, which was then visualised using iTOL software. The different lineages in the *X. vasicola* population are represented by the inner colour ring. The presence of BXO8 type (pink) and Xvv type LPS cassettes is indicated by the outermost arrow (sky blue color). (**B**). Comparison of BXO8 type LPS (pink) with Xvv NCPPB 795 (pink) and Xvv SAM119 (pink) (sky blue). The *etfA* and *metB* genes are represented by grey arrows, while IS elements are shown in red.

**Figure 4:**
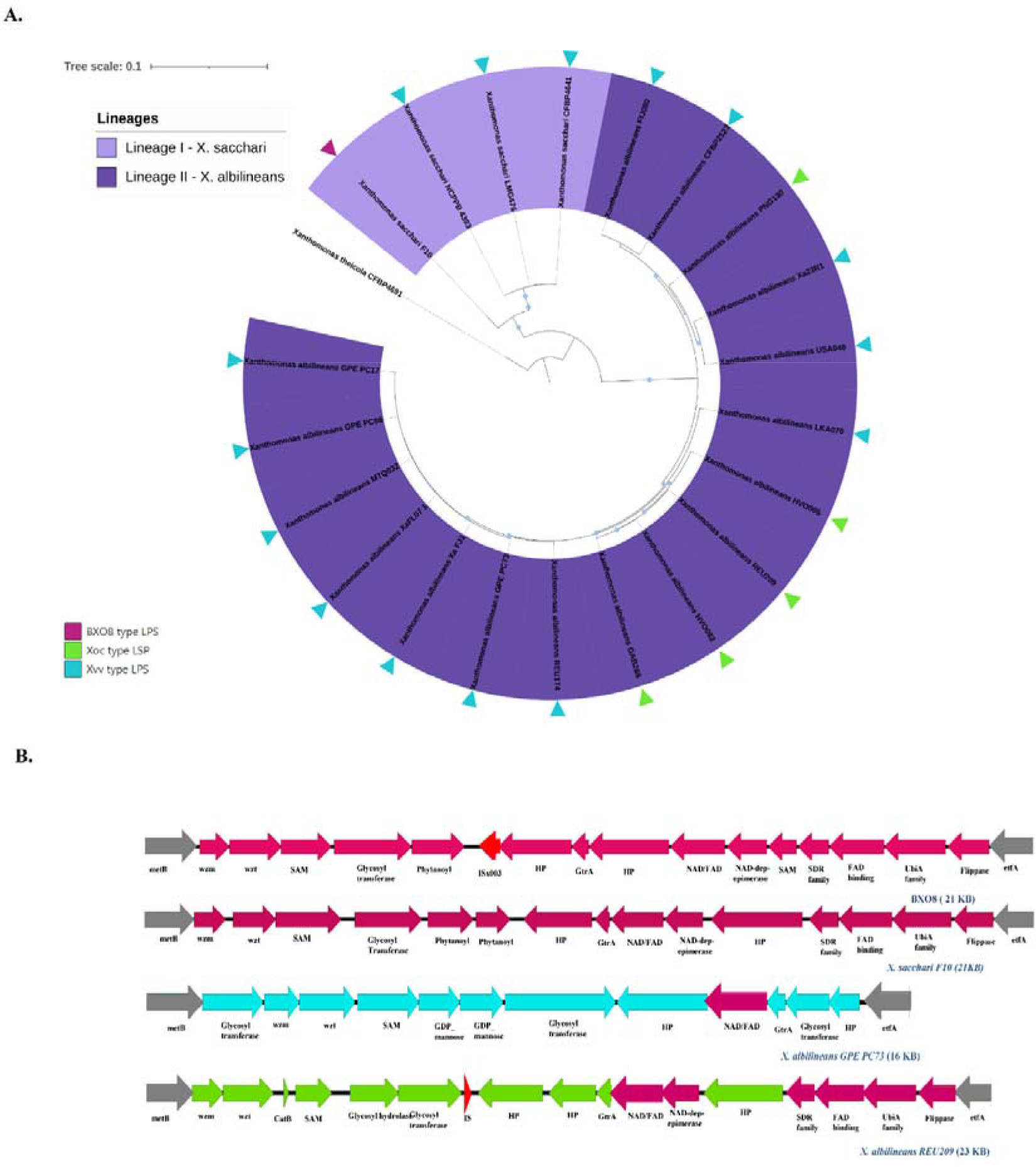
Phylogenetic tree of *X. albilineans* and *X. sacchari* populations. (**A**) Population study based on core genes and visualised with iTOL software, lineage-I (dark purple) and X. sacchari, lineage-II (light purple) (light purple). The presence of BXO8 type (pink colour), Xvv type (sky blue colour), and Xoc type (green colour) LPS cassettes in both populations is represented by the outermost arrow. (**B**). Comparison of LPS cassettes and genetic organisation in two populations with BXO8 type LPS. Pink represents BXO8 type LPS (BXO8 and X. sacchari F10), while sky blue and green represent Xvv (*X. albilineans* GPE PC73) and Xoc (*X. albilineans* REU209). The *etfA* and *metB* genes are represented by grey arrows, while IS elements are shown in red.

### Population mapping reveals that BXO8 has been replaced by newer types of LPS cassette(s) during their emergence

The availability of a large number of genome sequences from several *Xanthomonas* pathogens allowed us to understand the evolution of the LPS locus at the population level. In the case of the Xoo pathogen, which infects rice xylem, genome sequences of 406 strains are available in the public domain. While the BXO8 type gene cluster is found in 19 strains, the remaining 386 strains consist of the entirely distinct BXO1 type LPS cluster (figure 2A, B). The phylogenomic analysis reveals that the BXO1 type of cassette has overtaken a 386/406 Xoo strains (figure 2A). The BXO1 LPS cassette is 14 KB in size, has a 54% GC content, and encodes nine genes that have homology to genes involved in LPS biosynthesis and transport. In BXO1, the *wzm* and *wzt* genes are on the *etfA* side, whereas they are on the metB side in the BXO8 type LPS cassette. The presence of a large number of continuous transposes (IS5, IS256, IS30, and IS701) on the *etfA* side differentiates this LPS from BXO8. Even the *FkbM* family methyltransferase and acetyltransferase on the *metB* side are responsible for O-antigen biosynthesis of BXO1 type LPS, distinguishing it from BXO8 type LPS (figure 2B, supplementary figure 1).

Aside from Xoo, we have discovered BXO8 LPS cassette in *X*. *vasicola* (figure 1A), which is another pathogen with a large-scale genomic resource (n =113 genomes). As a result, we searched for the LPS locus diversity in the LPS of the Xvv population. The analysis revealed that only 19 strains have the BXO8 type of cassette, while 94 strains harbour a novel LPS cassette, which we name as novel Xvv type of LPS cassette (figure 3A and B). An Xvv type of cassette was reported in an earlier study while comparing variation at the LPS locus in *Xanthomonas* species that infect sugarcane. Interestingly, like in Xoo, the population level analysis reveals that in Xvv, the BXO8 type of LPS cassette was supplanted by a novel Xvv type LPS cassette (figure 3A). The new cassette has a size of 23 KB, a GC content of 58.5%, and the *wzm* and *wzt* marker gene do not match with the marker genes of BXO8 type LPS, making it novel LPS. The presence of glycosyltransferase just before the *wzm* and *wzt* marker genes in the *metB* flanking region is completely new. Aside from that, two GDP-mannose 4, 6-dehydratases and an IS3 family transposase are noticeable. This cassette contains a relatively new class I SAM-dependent methyltransferase. The other half of the *etfA* flanking region contains ORFs that are similar to those found in the BXO8 LPS cassette, resulting in Xvv LPS being a diversified BXO8 cassette (figure 3B, supplementary figure 2).

While Xoo and Xvv are from clade II, clade I consists of *X. sacchari*, which is another pathogen of sugarcane infecting xylem (figure 1A). Despite the fact that there are fewer genome sequences available for Xsac (n = 4 genomes) than for Xoo (n = 406 genomes) and Xvv (n = 113 genomes), we discovered that only one strain i.e. *X. sacchari* F10, which forms the basal branch of the phylogenetic tree (figure 4A), has the BXO8 type of LPS cassette. While the other three strains have a new LPS cassette that is interestingly homologous to the Xvv type of LPS cassette (figure 4A and B). It appears that during the evolution of Xoo and Xvv pathogens from clade II and Xsac from clade I, selection of novel LPS cassettes rather than the BXO8 type of LPS occurred.

### The emergence of *X. oryzae* variant lineages infecting parenchyma is associated with the acquisition of a novel LPS cassette from its relative *X. citri pv. citri,* which infects the parenchyma of citrus plants

We used a total of 22 genomes of Xoc (n = 19 genomes) and Xol (n = 3 genomes) with the Xoo (n = 405 genomes) population that are available at NCBI (https://www.ncbi.nlm.nih.gov/).Interestinly, Xoc and Xol fall into lineage II only and are not found in any other lineages (figure 5A). Amongst seven lineages, it is the only lineage consisting solely of pathovars that infect specifically parenchymal tissues (figure 5A). A genomic analysis of the LPS locus in Xoc and Xol revealed that these pathovars harbour a distinct type of LPS as compared to xylem-infecting LPS (BXO1 and BXO8). In Xoc type LPS, seven genes towards the *metB* side were identified as distinct. The Xoc locus is 22 KB and has 58% GC content, making it larger than BXO1 and BXO8, while having the highest GC content. The *catB-O* acetyltransferase, which is present only in Xoc LPS, is a chloramphenicol resistance effector in bacteria. Xoc has a hybrid LPS cluster; half of this gene cluster is homologous to the LPS cluster of BXO8, while the other half is very distinct (figure 5B, supplementary figure 1). The genes from the *etfA* side of Xoc LPS cluster such as Flippase-like domain, *UbiA*, *FAD*-binding, *NAD/FAD* oxidoreductase, and the *GtrA* family, were homologous to those from the BXO8 LPS cluster. Further, *SDR* family oxidoreductase, two different glycosyltransferases, and considerable numbers of hypothetical proteins are present in Xoc type LPS that are absent in other cassettes which are absent in rice xylem infecting LPS. The other side of *metB* was essentially different from the BXO8 LPS cluster, inferring it a chimeric LPS gene cluster and utterly different from the BXO1 type of cluster (supplementary figure 1). This chimeric LPS cluster of Xoc might be responsible for the origin of tissue-specific pathogenicity of Xoc and helping it evolve into a parenchyma-specific pathovar. Interestingly, the LPS cassette in Xol is of 25 KB with 57.5% GC content and contain same marker genes (*wzm* and *wzt*) as in the Xoc type (figure 5B). Numerous glycosyltransferases and hydrolases, however, are replaced, and hypothetical proteins and IS elements are introduced (figure 5B). This addition of five hypothetical proteins with unknown functions may aid in the infection of grasses and the evolution of Xol as a parenchyma pathovar capable of jumping from grasses to paddies. Despite the presence of Xoc type marker genes (*wzm* and *wzt*) in the LPS of Xol strains, the addition of six genes between the cassettes demonstrates that HGT occurs within the cassette to allow it to survive in two distinct hosts. All the strains of *X. oryzae* that infect parenchyma strains have a Xoc type of LPS cassette, whereas BXO1 and BXO8 cassettes are specifically present in Xoo strains that infect xylem tissue.

**Figure 5:**
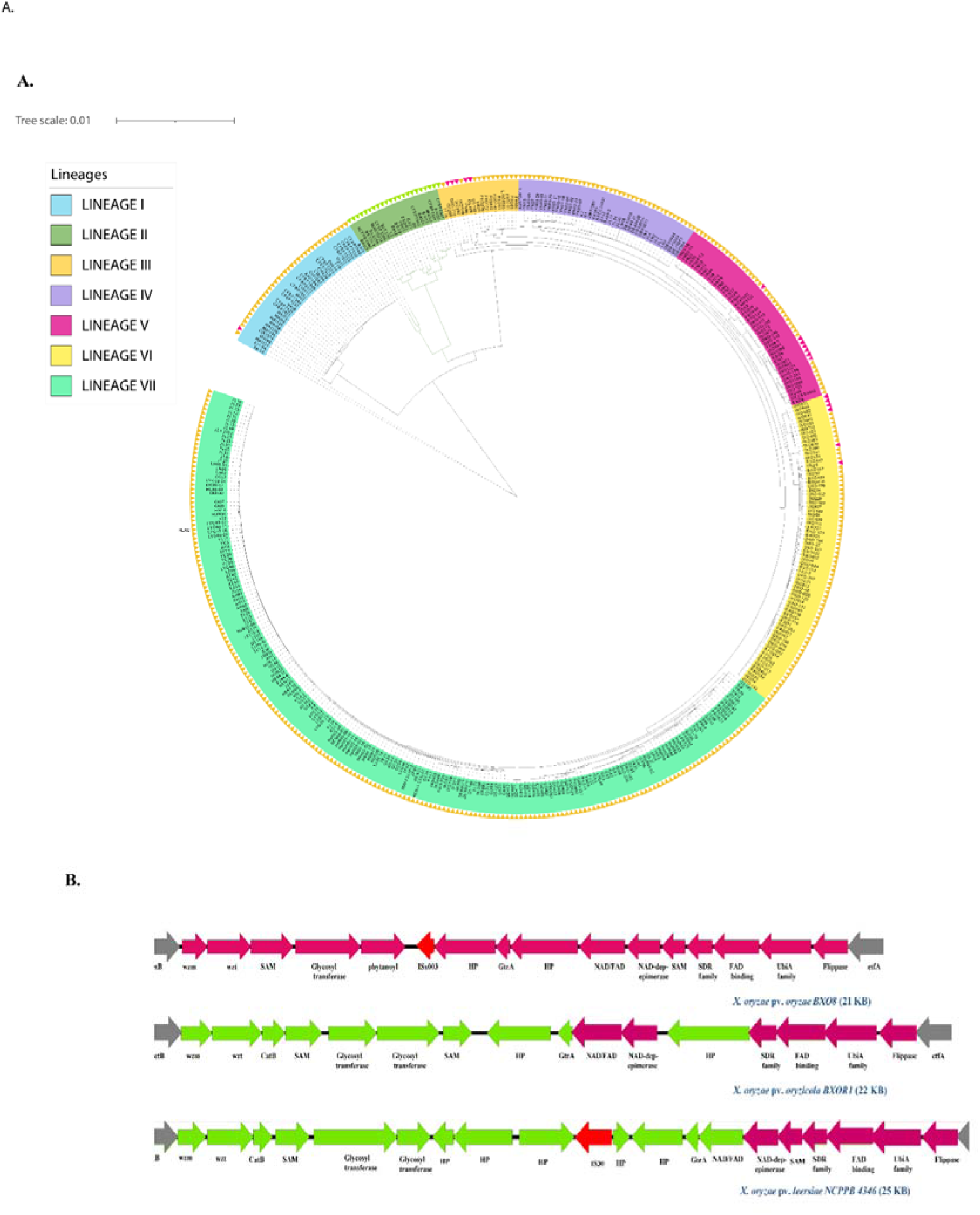
Distribution of Xoc and Xol lineages and LPS cassettes in the Xoo population. (**A**) PhyML was used to reconstruct the core-gene tree, which was then visualised using iTOL software. Lineages are represented by different node colours. The green node represents the Xoc and Xol population, with * denoting parenchyma pathovar and #Xoo denoting Xoo-xylem pathovar. The outermost arrows indicate the presence of LPS cassettes (yellow-BXO1, green-Xoc type, pink-BXO8) in each strain. The number of substitutions per site is indicated by the scale bar. (**B**) Comparison of Xoc type LPS (*X. oryzae* pv. *oryzicola* BXOR1 and *X. oryzae* pv. *leersia* NCPPB 4346) and BXO8 type LPS cassette the *etfA* and *metB* genes are represented by grey arrows, while IS elements are shown in red.

NCBI BLAST results indicate that the Xoc type is found in the genomes of strains of *Xanthomonas* pathogens, *X. citri pv. citri* (Xcc), *X. fragariae* YL19, and *X. arboricola pv. corylina* A7, which infect the parenchyma of citrus, strawberry, and hazelnut plants, respectively. Further, we mapped the distribution of the Xoc type of cassette in the publicly available 200 genome sequences of the Xcc population (figure 6B). We discovered that 14 strains that are A* and Aw pathotypes, have the Xoc type of LPS cassette arrangement with 100% coverage and more than 90% percent identity with Xoc LPS (figure 6B). It is pertinent to note that the Xcc causes citrus bacterial canker and is found to exist as three distinct pathotypes: A, A*, and Aw, of which has a unique host range and host response variations (Webster, Bogema, & Chapman, 2020). Pathotype A has the broadest host range, whereas pathotypes A* and Aw have a narrower host range that infects key lime (*Citrus aurantifolia*), alemow (*Citrus macrophylla),* and produces a hypersensitive response on grapefruit (Jalan et al., 2013; Rybak, Minsavage, Stall, & Jones, 2009). The evolutionary study suggests that both A* and Aw changed from a wide host range to a narrow host range over time. Whereas *X. fragariae* YL19 has a limited host range, it primarily infects strawberry varieties, causing crown infection pockets and angular leaf spots. Meanwhile, *X. arboricola* pv*. corylina* A7 infects hazels. *X. arboricola* pv*. corylina* A7 has over 90% query coverage and 54% identity with Xoc LPS, whereas *X. fragariae* has 100% query coverage and 94% identity with Xoc LPS (figure 6B, supplementary figure 3).

**Figure 6:**
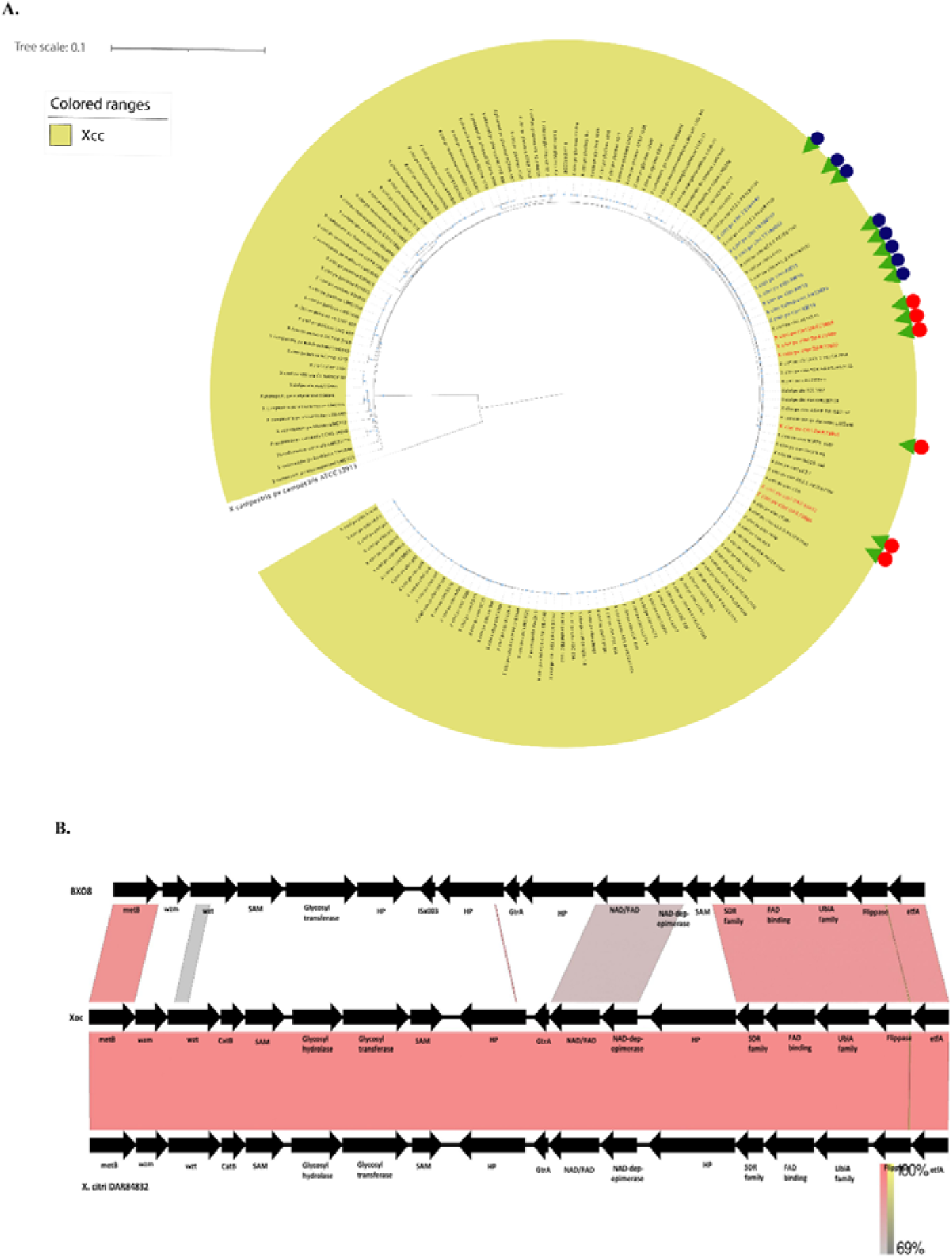
Distribution of the Xoc type LPS in Xcc population. (**A**) PhyML was used to reconstruct the core-gene tree, which was then visualised using iTOL software. The outermost symbols represent different pathotypes, with green colour arrows representing strains with Xoc type LPS with A* pathotype (red colour) and Aw pathotype (blue colour) (blue color). (**B**). Easyfig analysis: Xoc type LPS and BXO8 LPS cluster comparison with *X. citri* pv. *citri* DAR84832. The arrows indicate gene location, and the shaded lines indicate the degree of homology between pairs of genes in two LPS cassettes.

Hence, the presence of the Xoc type of LPS cassette in these pathotypes indicates that this emerging Xoc type is undergoing continuous evolution in *Xanthomonas*, assisting bacteria in evolving as parenchyma pathovars and restricting them becoming host-specific pathovars as well. The presence of the Xoc type of LPS cassette in diverse *Xanthomonas* species infecting the parenchyma of diverse hosts indicates a potential role of this type of cassette in parallel evolution. It is possible that HGT’s of Xoc LPS cassette in a Xoo strain played a critical role in the emergence of the *X. oryzae* pathovar capable of infecting the rice parenchyma. The other pathovar of *X. oryzae*, i.e., Xol, also infects the parenchyma of *Leersia hexandra* grass and forms a monophyletic lineage with Xoc. Unsurprisingly, Xol strains also comprise Xoc-type LPS cassette, corroborated our observation.

### Distinct LPS cassettes support diversified tissue specificities in sugarcane infecting Xalb pathogen

Xalb, another significant *Xanthomonas* pathogen that causes leaf scald, a lethal sugarcane disease, is also found in clade I (figure 1A). The NCBI public domain contains 17 genome sequences, as a result we scanned the LPS locus in these strains and discovered that Xalb is dominated by the Xvv type of LPS cassette. Twelve of the seventeen strains contain an LPS cassette of the Xvv type, while the remaining five contain an LPS cassette of the Xoc type (figure 4A). The Xvv type of LPS in Xalb has query coverage of more than 50%, whereas the *wzm* maker gene has 100% query coverage with an Xvv type LPS cassette, making it more similar to sugarcane infecting like the Xvv type LPS cassette. In comparison to the Xvv type of *X. vasicola,* the Xalb Xvv type LPS cassette is smaller in size (16 KB) with 58.7% GC content. The arrangement of genes towards *metB* is similar to that of the Xvv type where there is a reduction of genes like the *SDR* family, *UbiA* family, and *Flippase* on the *etfA* side. The presence of hypothetical genes, glycosyltransferas, and *GtrA* family proteins flanking the *etfA* gene is unique to this cassette (figure 4B). In the case of Xoc type LPS in Xalb, it is 23 KB in size (55% GC), whereas the pseudogenization of Xoc specific *catB-O* acetyltransferase towards the *metB* side is noticeable. Along with this, there is the presence of two glycosyltransferases and three hypothetical proteins with unknown functions on the *etfA* side (figure 4B). Infection and multiplication of Xalb bacteria in sugarcane xylem along with parenchyma tissues have been reported recently. Hence, we can conclude that in order to successfully invade both vascular and non-vascular sugarcane tissue, it appears that two distinct LPS types are required in the Xalb population.

## Discussion

Large-scale variation in pathogens can be expected because of the enormous diversification of the hosts they infect. Further, with the advancement of third-generation sequencing, it becomes easier to study complete genome sequences and find their origin and evolution through comparative and population genomics. The large-scale variations in LPS clusters of *Xanthomonas* at strain, pathovar, and species-level indicate that plant pathogenic bacteria are under intense selection to vary their *lps* gene cluster, and earlier mutation studies suggest an essential role for LPS in plant-pathogen interactions and the virulence process (Petrocelli, Tondo, Daurelio, & Orellano, 2012).

Our deep phylogenomics of diverse *Xanthomonas* species reveals lineages associated with different types of LPS in accordance with their host and tissue specificity (figure 7). Population study of rice pathogens suggests it was originally a xylem pathogen, as evident by multiple Xoo pathovar lineages versus a single parenchyma lineage comprising of Xoc and Xol sandwiched between xylem Xoo lineages. As a result, parenchyma pathovars could be variant lineages of the xylem pathovars in the *X. oryzae* population (Figure 2A). Further, the presence of the Xoc type LPS cassette in the narrow host-range Xcc population implies their significance in invading the parenchyma tissues of their respective hosts. To support our finding, we have also found that pathogens that infect the parenchyma, such as *X. fragariae* and *X. arboricola* pv. *corylina* also have Xoc type LPS cassettes.

**Figure 7:**
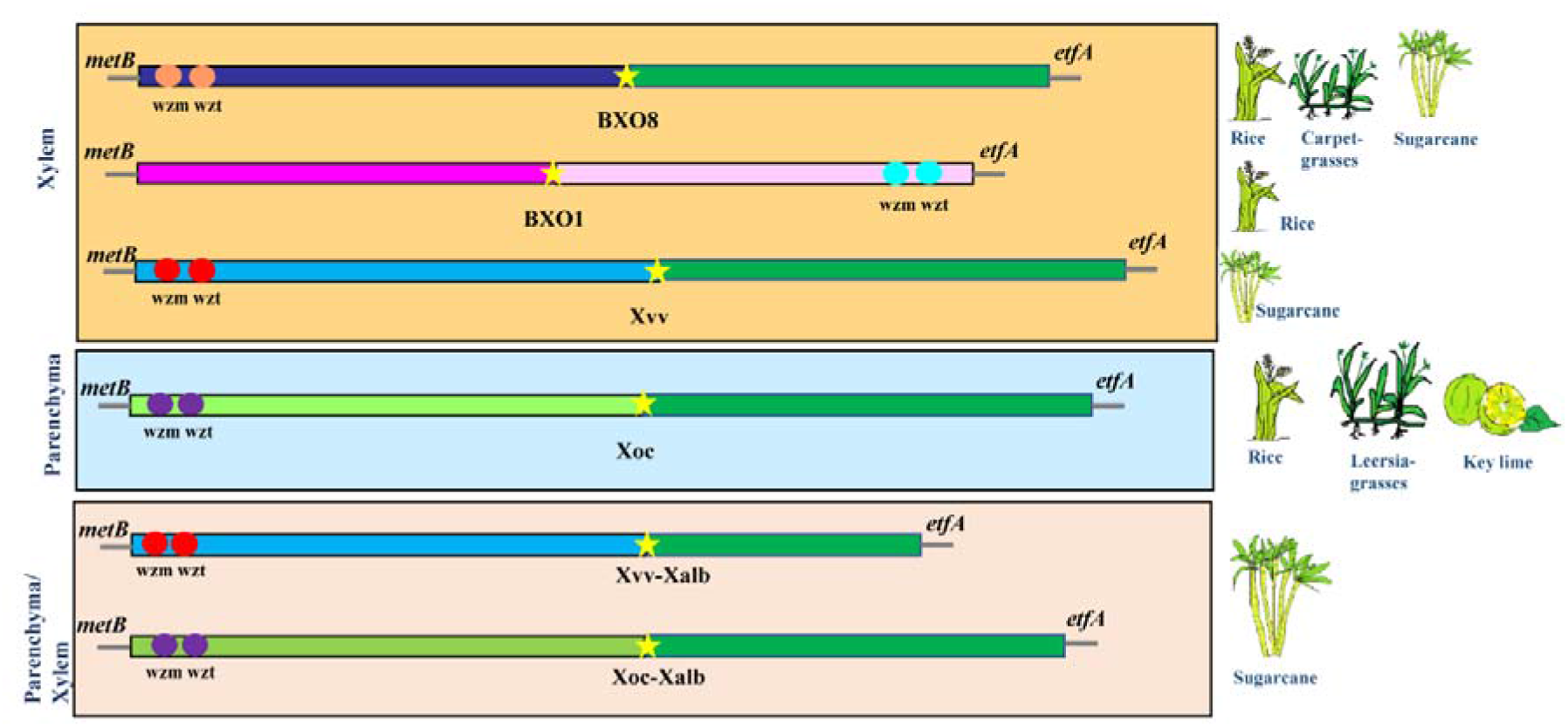
Schematic view representing the major type of LPS cassettes in *Xanthomonas* population. On the left are names of different tissues (indicating color boxes), and on the right are different hosts to which each type of LPS is linked. Each LPS cassette has *metB* and *etfA* flanking genes. The *wzm* and *wzt* marker genes are shown next to each other in the same colour scheme, indicating homology. The star signifies IS elements and hypothetical genes. In present study *metB* locus region of each cassette is found to be the most hyper-variable, responsible for virulence and transport. The other biosynthesis locus is on *etfA* region and contains genes that are similar to the ancestral one (BXO1 LPS is an exception).

The presence of BXO8 type cassettes in xylem pathogens from distinct clades, such as *X. oryzae* infecting rice, *X. axonopodis,* which infects grasses, and sugarcane pathogens *X. vasicola* and *X. sacchari*, suggests that it was an ancestral cassette. Interestingly, in xylem invaders, BXO8 type LPS is supplanted by BXO1 in rice pathogens and Xvv type LPS in sugarcane pathogens. Our study illustrates that genes at the *metB* locus of BXO8 are at systemic hyper-variation due to presence of multiple IS elements, leading to distinct glycosyltransferases, methyltransferases, and hypothetical proteins. In this context, distinct LPS cassettes fit the bill as a dominant emerging lineage leads to successful invasion and adaptation in sugarcane and rice plants.

Peculiarly, *X. albilineans* is a lethal sugarcane pathogen with reduced genome invading both xylem and parenchyma (Pieretti et al., 2015). Our genomic investigation reveals Xalb to have two different LPS cassettes: Xvv and Xoc types, found to be specific to xylem invading sugarcane pathogens and parenchyma invaders in the present study. Having two different types of LPS cassettes without any pathovar or tissue specific lineage suggests that these pathovars have evolved their LPS cassettes to survive in both xylem and parenchyma tissues. This also supports our conclusion that LPS not only plays a role in pathovar specific lineage diversification, but also in pathogen adaptation to different hosts and tissue specificity. Unlike rice pathogens, Xalb lacks tissue-specific lineages warranting its population level genomic investigation.

Both the xylem and parenchyma represent distinct niches in the plant (An et al., 2020). In the xylem, it is expected that defence responses are lower compared to the parenchyma. Parenchyma pathogens may have to actively deal with antimicrobial and other defence responses of parenchyma cells. LPS may act as barrier for antimicrobial peptides and molecules (Galloway & Raetz, 1990). In *Burkholderia cenocepacia*, deletion mutants in the LPS gene cluster were found to be resistant to tailocin (a phage tail-like bacteriocin) (Yao et al., 2017). Because LPS is a phage receptor, xylem pathovars are likely to have a different type of LPS cassette than parenchyma pathogens. Simultaneously, the number and diversity of phages in the xylem may be higher than in the parenchyma due to the continuity of water flow. While *Xanthomonas* pathogens that infect the xylem of grasses have acquired Xoc type cassette which led to the emergence and success of *X. oryz*ae as a parenchyma pathogen may have coincided with the acquisition of LPS cassettes, which are also present in strains of Xol. The deep genomic investigation has provided detailed insights into changes in large-scale variations mediated by horizontal gene transfer in diverse *Xanthomonas* pathogens that infect the xylem and parenchyma of grasses, sugarcane, and citrus plants. Further genetic, cellular, and functional studies are warranted to establish the role of variant LPS cassettes in conferring success on these tissue-specific pathovars and their predominant lineages.

## Materials and Methods

### Genome procurement from the public repository

A total of 427 (Xoo), 215 (Xcc), 114 (Xvv), 63 (Xtl), 17 (Xalb), 4 (Xsac), and 5 (Xaxn) genomes were used in this study. All *Xanthomonas* genomes [in 427 genomes, Xoo (n = 405), Xoc (n = 19), and Xol (n = 3)] are publicly available (https://www.ncbi.nlm.nih.gov/) and were used in this study.

### Pan-genome analysis

Roary v3.12.0 (Page et al., 2015) was used to perform pan-genome analysis using .gff files generated by Prokka v1.13.3 (Seemann, 2014) as input. As we were analysing genomes from the same species, the cut-off used was 96%, which is set by default.

### Phylogenetic analysis

A phylogenetic tree analysis was performed using MEGA7 (Kumar, Stecher, & Tamura, 2016) by the neighbor-joining method. A core genome tree of genomes was constructed using PhyML (Guindon et al., 2010). For example, the core genome alignment was generated using Roary v3.11.2 (Page et al., 2015), and it was converted to phylip format using SeaView v4.4.2-1 (Gouy, Guindon, & Gascuel, 2010). Then, the Newick tree file was obtained by using PhyML and then visualized using iTOL (Letunic & Bork, 2021).

### Comparative study-LPS cassette analysis

All the LPS cassettes flanked by two conserved sequences on both sides of the cassette were extracted from genomes by doing NCBI BLAST (Johnson et al., 2008) of marker *wzm* (1 KB) and *wzt* (1 KB) genes. Firstly, the query coverage and percent identity (more than 50%) of the *wzm* gene are used to extract the cassettes from all genomes (both protein and nucleotide sequence), and then a complete cassette (both protein and nucleotide sequence) is BLAST against each LPS type of LPS cassette on the basis of query coverage and percent identity (more than 50%) of the complete cassette for comparisons. With the help of an easy genome comparison tool-Easyfig v2.2 (Sullivan, Petty, & Beatson, 2011) (https://mjsull.github.io/Easyfig/), we performed BLAST and aligned all four complete LPS cassettes with each other. We have also used the Artemis tool for annotation and visualization of sequences of cassettes (http://sanger-pathogens.github.io/Artemis/Artemis/) (Rutherford et al., 2000).

## Abbreviations

HGT: horizontal gene transfer
LPS: lipopolysaccharide
Xoo: Xanthomonas oryzae pv. oryzae
Xoc: Xanthomonas oryzae pv. oryzicola
Xol: Xanthomonas oryzae pv. leersia
Xcc: Xanthomonas citri pv. citri
Xvv: Xanthomonas vasicola pv. vasculorum
Xalb: Xanthomonas albilineans
Xsac: Xanthomonas sacchari
Xaxn: Xanthomonas axonopodis
QC: query coverage
Per.ID: percent identity
PAMP: pathogen-associated molecular pattern

## Authors contributions

AS performed the data analysis and drafted the manuscript with inputs from KB, SK, and PBP. PBP conceived and participated in designing the study and finalizing the manuscript.

## Acknowledgment

We acknowledge the funding through NBRI-IMTECH-MLP48 and the CSIR fellowship.

## Transparency declaration

The authors declare that there are no conflicts of interest.

## Supplementary material

**Supplementary figure 1:** LPS synteny comparison of BXO1, BXO8 and Xoc type LPS cassette with Easyfig and the BLASTn algorithm. Arrows represent the location of genes and shaded lines reflect the degree of homology between pairs of genes in two LPS cassettes.

**Supplementary figure 2:** LPS synteny comparison with Easyfig and the BLASTn algorithm. Arrows represent the location of genes and shaded lines reflect the degree of homology between pairs of genes in two LPS cassettes. *X. sacchari* CFBP 4641 and *X. albilineans* CFBP 2523 shows more homology with Xvv type of LPS than BXO8.

**Supplementary figure 3:** LPS synteny comparison with Easyfig and the BLASTn algorithm. Arrows represent the location of genes and shaded lines reflects the degree of homology between pairs of genes in two LPS cassettes. Gene cluster in Xcc strains (DAR84832, AW13), *X. fragariae* YL19, *X. arboricola* pv. *corylina* A7 LPS showing homology with Xoc type LPS cassette (green color).

**Supplementary table 1:** Metadata of *Xanthomonas* strains used in present study.

## Notes

### Competing Interest Statement

The authors have declared no competing interest.

