## supplementary table 1 for "Systematic hyper-variation and evolution at a lipopolysaccharide locus in the population of *Xanthomonas* species that infect rice and sugarcane"

**Supplementary table 1**: Metadata of Xanthomonas strains used in present study.

| SPECIES | *wzm* gene QC% (nt) | *wzm gene* per.ID % (nt) | *wzt* gene QC% (nt) | *wzt* gene per.ID % (nt) | LPS TYPE | LPS per.ID % (nt) | SIZE (MB) | LEVEL | GC% | COMMON HOST | ASSEMBLY ID |
|---|---|---|---|---|---|---|---|---|---|---|---|
| X.oryzae pv. oryzae X11-5A | 100 | 99.88 | 100 | 100 | BXO1 | 99.87 | 4.95 | COMPLETE | 64 | Rice | GCA_001277045.1 |
| X.oryzae pv. oryzicola X8-1A | 100 | 66.33 | 100 | 68.88 | Xoc | 99.96 | 4.59 | DRAFT | 64 | Rice | GCA_000212775.2 |
| X.oryzae pv. oryzae MAI134 | 100 | 98.47 | 100 | 100 | BXO8 | 99.62 | 4.73 | COMPLETE | 63.9 | Rice | GCA_002850175.1 |
| X.oryzae pv. oryzae MAI99 | 100 | 99.88 | 100 | 100 | BXO1 | 99.87 | 4.7 | COMPLETE | 63.9 | Rice | GCA_002850215.1 |
| X.oryzae pv. oryzae CIX2374 | 100 | 99.88 | 100 | 100 | BXO1 | 99.87 | 4.73 | DRAFT | 63.9 | Rice | GCA_004322815.1 |
| X.oryzae pv. oryzae MAI145 | 100 | 99.88 | 100 | 100 | BXO1 | 99.87 | 4.7 | COMPLETE | 63.9 | Rice | GCA_002850075.1 |
| X.oryzae pv. oryzae MAI106 | 100 | 99.88 | 100 | 100 | BXO1 | 99.87 | 4.71 | COMPLETE | 63.9 | Rice | GCA_002850135.1 |
| X.oryzae pv. oryzae MAI129 | 100 | 99.88 | 100 | 100 | BXO1 | 99.87 | 4.7 | COMPLETE | 63.9 | Rice | GCA_002850155.1 |
| X.oryzae pv. oryzae MAI129 | 100 | 99.88 | 100 | 100 | BXO1 | 99.87 | 4.7 | COMPLETE | 63.9 | Rice | GCA_002850155.1 |
| X.oryzae pv. oryzae MAI73 | 100 | 99.88 | 100 | 100 | BXO1 | 99.87 | 4.7 | COMPLETE | 63.9 | Rice | GCA_002850075.1 |
| X.oryzae pv. oryzae MAI95 | 100 | 99.88 | 100 | 100 | BXO1 | 99.87 | 4.7 | COMPLETE | 63.9 | Rice | GCA_002850195.1 |
| X.oryzae pv. oryzae CFBP1951 | 100 | 99.88 | 100 | 100 | BXO1 | 99.87 | 4.7 | COMPLETE | 63.9 | Rice | GCA_004355445.1 |
| X.oryzae pv. oryzae CFBP7325 | 100 | 99.88 | 100 | 100 | BXO1 | 99.87 | 4.73 | COMPLETE | 63.9 | Rice | GCA_004355385.1 |
| X.oryzae pv. oryzae NAI8 | 100 | 99.88 | 100 | 100 | BXO1 | 99.87 | 4.34 | COMPLETE | 64.1 | Rice | GCA_000511585.1 |
| X.oryzae pv. oryzae CFBP7337 | 100 | 99.88 | 100 | 100 | BXO1 | 99.87 | 4.73 | COMPLETE | 63.9 | Rice | GCA_004355485.1 |
| X.oryzae pv. oryzae CFBP1949 | 100 | 99.88 | 100 | 100 | BXO1 | 99.87 | 4.74 | COMPLETE | 63.9 | Rice | GCA_004355605.1 |
| X.oryzae pv. oryzae CFBP1952 | 100 | 99.88 | 100 | 100 | BXO1 | 99.87 | 4.69 | COMPLETE | 63.9 | Rice | GCA_004355585.1 |
| X.oryzae pv. oryzae CFBP8172 | 100 | 99.88 | 100 | 100 | BXO1 | 99.87 | 4.66 | COMPLETE | 63.9 | Rice | GCA_004355345.1 |
| X.oryzae pv. oryzae CIX611 | 100 | 99.88 | 100 | 100 | BXO1 | 99.87 | 4.68 | COMPLETE | 63.9 | Rice | GCA_004299395.1 |
| X.oryzae pv. oryzae AXO1947 | 100 | 99.88 | 100 | 100 | BXO1 | 99.87 | 4.67 | COMPLETE | 63.9 | Rice | GCA_001466505.1 |
| X.oryzae pv. oryzae CFBP1948 | 100 | 99.88 | 100 | 100 | BXO1 | 99.87 | 4.67 | COMPLETE | 63.9 | Rice | GCA_004355465.1 |
| X.oryzae pv. oryzae CIX1041 | 100 | 99.88 | 100 | 100 | BXO1 | 99.87 | 5.01 | DRAFT | 63.8 | Rice | GCA_004299365.1 |
| X.oryzae pv. oryzae CIX298 | 100 | 99.88 | 100 | 100 | BXO1 | 99.87 | 4.75 | COMPLETE | 63.9 | Rice | GCA_004322835.1 |
| X.oryzae pv. oryzae CFBP7322 | 100 | 99.88 | 100 | 100 | BXO1 | 99.87 | 4.71 | COMPLETE | 63.9 | Rice | GCA_004355525.1 |
| X.oryzae pv. oryzae CIX629 | 100 | 99.88 | 100 | 100 | BXO1 | 99.87 | 4.74 | DRAFT | 63.9 | Rice | GCA_004321555.1 |
| X.oryzae pv. oryzae DAK16 | 100 | 99.88 | 100 | 100 | BXO1 | 99.87 | 4.72 | DRAFT | 63.9 | Rice | GCA_004355325.1 |
| X.oryzae pv. oryzae CFBP7324 | 100 | 99.88 | 100 | 100 | BXO1 | 99.87 | 4.72 | COMPLETE | 63.9 | Rice | GCA_004355505.1 |
| X.oryzae pv. oryzae T19 | 100 | 99.88 | 100 | 100 | BXO1 | 99.87 | 4.72 | COMPLETE | 63.9 | Rice | GCA_004355305.1 |
| X.oryzae pv. oryzae Ug11 | 100 | 99.88 | 100 | 100 | BXO1 | 99.87 | 4.72 | COMPLETE | 63.9 | Rice | GCA_004355285.1 |
| X.oryzae pv. oryzae CFBP7321 | 100 | 99.88 | 100 | 100 | BXO1 | 99.87 | 4.72 | COMPLETE | 63.9 | Rice | GCA_004355425.1 |
| X.oryzae pv. oryzae CFBP7319 | 100 | 99.88 | 100 | 100 | BXO1 | 99.87 | 4.75 | COMPLETE | 63.9 | Rice | GCA_004355565.1 |
| X.oryzae pv. oryzae CFBP7323 | 100 | 99.88 | 100 | 100 | BXO1 | 99.87 | 4.73 | COMPLETE | 63.9 | Rice | GCA_004355405.1 |
| X.oryzae pv. oryzae CFBP7320 | 100 | 99.88 | 100 | 100 | BXO1 | 99.87 | 4.72 | COMPLETE | 63.9 | Rice | GCA_004355545.1 |
| X.oryzae pv. oryzae BAI3 | 100 | 99.88 | 100 | 100 | BXO1 | 99.87 | 4.72 | COMPLETE | 63.9 | Rice | GCA_003031385.1 |
| X.oryzae pv. oryzae CFBP7340 | 100 | 99.88 | 100 | 100 | BXO1 | 99.87 | 4.74 | COMPLETE | 63.9 | Rice | GCA_004355365.1 |
| X.oryzae pv. leersia BAI23 | 100 | 66.55 | 100 | 68.86 | Xoc | 94.52 | 4.75 | COMPLETE | 64 | Rice | GCA_004319505.1 |
| X.oryzae pv. leersia BB156-2 | 100 | 66.55 | 100 | 68.86 | Xoc | 94.52 | 5.07 | COMPLETE | 63.8 | Rice | GCA_004319465.1 |
| X.oryzae pv. leersia NCPPB4346 | 100 | 66.55 | 100 | 68.86 | Xoc | 94.52 | 4.95 | COMPLETE | 63.9 | Rice | GCA_001276975.2 |
| X.oryzae pv. oryzicola BLS256 | 100 | 99.76 | 100 | 99.64 | Xoc | 99.86 | 4.83 | COMPLETE | 64.1 | Rice | GCA_000168315.3 |
| X.oryzae pv. oryzicola CFBP2286 | 100 | 99.64 | 100 | 99.64 | Xoc | 99.86 | 5.01 | COMPLETE | 63.9 | Rice | GCA_001042735.1 |
| X.oryzae pv. oryzicola WHRI 5234 | 100 | 99.64 | 100 | 99.64 | Xoc | 99.86 | 4.69 | DRAFT | 64.1 | Rice | GCA_003064145.1 |
| X.oryzae pv. oryzicola BLS279 | 100 | 99.76 | 100 | 99.64 | Xoc | 99.86 | 4.83 | COMPLETE | 64.1 | Rice | GCA_001042775.1 |
| X.oryzae pv. oryzicola L8 | 100 | 99.64 | 100 | 99.64 | Xoc | 99.86 | 4.8 | COMPLETE | 64.1 | Rice | GCA_001042855.1 |
| X.oryzae pv. oryzicola B8-12 | 100 | 99.64 | 100 | 99.64 | Xoc | 99.86 | 4.79 | COMPLETE | 64.1 | Rice | GCA_001042745.1 |
| X.oryzae pv. oryzicola B812 | 100 | 99.64 | 100 | 99.64 | Xoc | 99.86 | 4.79 | COMPLETE | 64.1 | Rice | GCA_001042745.1 |
| X.oryzae pv. oryzicola BXOR1 | 100 | 99.64 | 100 | 99.64 | Xoc | 99.86 | 4.69 | COMPLETE | 64.1 | Rice | GCA_001042795.1 |
| X.oryzae pv. oryzicola YM15 | 100 | 99.88 | 100 | 99.64 | Xoc | 99.86 | 4.43 | COMPLETE | 64.1 | Rice | GCA_001021915.1 |
| X.oryzae pv. oryzicola 9 | 100 | 99.64 | 100 | 99.64 | Xoc | 99.86 | 4.83 | COMPLETE | 64 | Rice | GCF_009660205.1 |
| X.oryzae pv. oryzicola GX01 | 100 | 99.88 | 100 | 99.64 | Xoc | 99.86 | 4.85 | COMPLETE | 63.97 | Rice | GCA_020084945.1 |
| X.oryzae pv. oryzicola BAI21 | 100 | 99.64 | 100 | 99.64 | Xoc | 99.86 | 4.26 | DRAFT | 64.2 | Rice | GCA_002189465.1 |
| X.oryzae pv. oryzicola CFBP7342 | 100 | 99.64 | 100 | 99.64 | Xoc | 99.86 | 5.08 | COMPLETE | 64 | Rice | GCA_000940825.1 |
| X.oryzae pv. oryzicola BAI15 | 100 | 99.64 | 100 | 99.64 | Xoc | 99.86 | 4.29 | DRAFT | 64 | Rice | GCA_002189395.1 |
| X.oryzae pv. oryzicola CFBP7331 | 100 | 99.64 | 100 | 99.64 | Xoc | 99.86 | 5.02 | COMPLETE | 64 | Rice | GCA_001042815.1 |
| X.oryzae pv. oryzicola MAI10 | 100 | 99.76 | 100 | 99.64 | Xoc | 99.86 | 4.42 | DRAFT | 64 | Rice | GCA_002850135.1 |
| X.oryzae pv. oryzicola BAI20 | 100 | 99.76 | 100 | 99.64 | Xoc | 99.86 | 4.47 | DRAFT | 64 | Rice | GCA_002189435.1 |
| X.oryzae pv. oryzicola CFBP7341 | 100 | 99.64 | 100 | 99.64 | Xoc | 99.86 | 5.01 | COMPLETE | 63.9 | Rice | GCA_000940825.1 |
| X.oryzae pv. oryzae FJ16 | 100 | 99.64 | 100 | 99.64 | Xoc | 96.45 | 4.625 | DRAFT | 63.7 | Rice | GCA_003297765.1 |
| X.oryzae pv. oryzae OS209 | 100 | 99.88 | 100 | 100 | BXO1 | 99.87 | 4.13 | DRAFT | 63.1 | Rice | GCA_003300615.1 |

| | | | | | | | | | | | |
|---|---|---|---|---|---|---|---|---|---|---|---|
| X.oryzae pv. oryzae DXO-216 | 100 | 98.6 | 100 | 100 | BXO8 | 99.62 | 4.267 | DRAFT | 63.9 | Rice | GCA_001927875.1 |
| X.oryzae pv. oryzae IXO599 | 100 | 98.6 | 100 | 100 | BXO8 | 99.62 | 4.306 | DRAFT | 63.9 | Rice | GCA_001928855.1 |
| X.oryzae pv. oryzae IXO597 | 100 | 98.6 | 100 | 100 | BXO8 | 99.62 | 4.28 | DRAFT | 63.9 | Rice | GCA_001928835.1 |
| X.oryzae pv. oryzae YN06 | 100 | 99.88 | 100 | 100 | BXO1 | 99.87 | 4.522 | DRAFT | 63.8 | Rice | GCA_003299625.1 |
| X.oryzae pv. oryzae OS109 | 100 | 98.6 | 100 | 100 | BXO8 | 99.62 | 4.117 | DRAFT | 63.1 | Rice | GCA_003301345.1 |
| X.oryzae pv. oryzae OS189 | 100 | 98.6 | 100 | 100 | BXO8 | 99.62 | 4.316 | DRAFT | 63.3 | Rice | GCA_003300685.1 |
| X.oryzae pv. oryzae YN30 | 100 | 99.88 | 100 | 100 | BXO1 | 99.87 | 4.358 | DRAFT | 63.4 | Rice | GCA_003299225.1 |
| X.oryzae pv. oryzae PXO236 | 100 | 99.88 | 100 | 100 | BXO1 | 99.87 | 4.969 | DRAFT | 63.7 | Rice | GCA_001746655.1 |
| X.oryzae pv. oryzae PXO112 | 100 | 99.88 | 100 | 100 | BXO1 | 99.87 | 4.654 | DRAFT | 63.7 | Rice | GCA_003294545.1 |
| X.oryzae pv. oryzae PXO145 | 100 | 99.88 | 100 | 100 | BXO1 | 99.87 | 5.04 | DRAFT | 63.7 | Rice | GCA_001746615.1 |
| X.oryzae pv. oryzae LMG9585 | 100 | 99.88 | 100 | 100 | BXO1 | 99.87 | 4.53 | DRAFT | 63.8 | Rice | GCA_001974485.1 |
| X.oryzae pv. oryzae PXO83 | 100 | 99.88 | 100 | 100 | BXO1 | 99.87 | 5.025 | DRAFT | 63.7 | Rice | GCA_001518895.1 |
| X.oryzae pv. oryzae PX086 | 100 | 99.88 | 100 | 100 | BXO1 | 99.87 | 5.02 | COMPLETE | 63.7 | Rice | GCA_003382915.1 |
| X.oryzae pv. oryzae PXO211 | 100 | 99.88 | 100 | 100 | BXO1 | 99.87 | 5.033 | DRAFT | 63.7 | Rice | GCA_001746635.1 |
| X.oryzae pv. oryzae PXO86 | 100 | 99.88 | 100 | 100 | BXO1 | 99.87 | 5.02 | COMPLETE | 63.7 | Rice | GCA_000948075.1 |
| X.oryzae pv. oryzae AUST2003 | 100 | 99.88 | 100 | 100 | BXO1 | 99.87 | 4.9 | COMPLETE | 63.7 | Rice | GCA_004355785.1 |
| X.oryzae pv. oryzae YN03-03 | 100 | 99.88 | 100 | 100 | BXO1 | 99.87 | 4.5 | DRAFT | 63.8 | Rice | GCA_003299955.1 |
| X.oryzae pv. oryzae YN15 | 100 | 99.88 | 100 | 100 | BXO1 | 99.87 | 4.4 | DRAFT | 63.6 | Rice | GCA_003299365.1 |
| X.oryzae pv. oryzae YN07 | 100 | 99.88 | 100 | 100 | BXO1 | 99.87 | 4.3 | DRAFT | 63.4 | Rice | GCA_003299875.1 |
| X.oryzae pv. oryzae YN03-09 | 100 | 99.88 | 100 | 100 | BXO1 | 99.87 | 4.2 | DRAFT | 63.3 | Rice | GCA_003299875.1 |
| X.oryzae pv. oryzae YN03-11 | 100 | 99.88 | 100 | 100 | BXO1 | 99.87 | 4.5 | DRAFT | 63.8 | Rice | GCA_003294365.1 |
| X.oryzae pv. oryzae YN03-03 | 100 | 99.88 | 100 | 100 | BXO1 | 99.87 | 4.5 | DRAFT | 63.8 | Rice | GCA_003299955.1 |
| X.oryzae pv. oryzae YN03-20 | 100 | 99.88 | 100 | 100 | BXO1 | 99.87 | 4.5 | DRAFT | 63.8 | Rice | GCA_003299745.1 |
| X.oryzae pv. oryzae YN03-15 | 100 | 99.88 | 100 | 100 | BXO1 | 99.87 | 4.5 | DRAFT | 63.8 | Rice | GCA_003294345.1 |
| X.oryzae pv. oryzae YN03-21 | 100 | 99.88 | 100 | 100 | BXO1 | 99.87 | 4.5 | DRAFT | 63.8 | Rice | GCA_003299665.1 |
| X.oryzae pv. oryzae YN03-18 | 100 | 99.88 | 100 | 100 | BXO1 | 99.87 | 4.5 | DRAFT | 63.8 | Rice | GCA_003294295.1 |
| X.oryzae pv. oryzae YN03-02 | 100 | 99.88 | 100 | 100 | BXO1 | 99.87 | 4.5 | DRAFT | 63.8 | Rice | GCA_003300065.1 |
| X.oryzae pv. oryzae YN03-04 | 100 | 99.88 | 100 | 100 | BXO1 | 99.87 | 4.5 | DRAFT | 63.8 | Rice | GCA_003294405.1 |
| X.oryzae pv. oryzae YN03-07 | 100 | 99.88 | 100 | 100 | BXO1 | 99.87 | 4.5 | DRAFT | 63.8 | Rice | GCA_003299885.1 |
| X.oryzae pv. oryzae YN03-24 | 100 | 99.88 | 100 | 100 | BXO1 | 99.87 | 4.5 | DRAFT | 63.8 | Rice | GCA_003311715.1 |
| X.oryzae pv. oryzae K2 | 100 | 99.88 | 100 | 100 | BXO1 | 99.87 | 4.9 | COMPLETE | 63.7 | Rice | GCA_011604745.1 |
| X.oryzae pv. oryzae JW11089 | 100 | 99.88 | 100 | 100 | BXO1 | 99.87 | 5 | COMPLETE | 63.7 | Rice | GCA_004355765.2 |
| X.oryzae pv. oryzae K2 | 100 | 99.88 | 100 | 100 | BXO1 | 99.87 | 4.9 | COMPLETE | 63.7 | Rice | GCA_011604745.1 |
| X.oryzae pv. oryzae KACC10331 | 100 | 99.88 | 100 | 100 | BXO1 | 99.87 | 4.9 | COMPLETE | 63.9 | Rice | GCA_000007385.1 |
| X.oryzae pv. oryzae KX085 | 100 | 99.88 | 100 | 100 | BXO1 | 99.87 | 4.9 | COMPLETE | 63.7 | Rice | GCA_004355845.1 |
| X.oryzae pv. oryzae BJ84-3 | 100 | 99.88 | 100 | 100 | BXO1 | 99.87 | 4.6 | DRAFT | 63.8 | Rice | GCA_003298095.1 |
| X.oryzae pv. oryzae BJ12 | 100 | 99.88 | 100 | 100 | BXO1 | 99.87 | 4.2 | DRAFT | 63.2 | Rice | GCA_003295625.1 |
| X.oryzae pv. oryzae JL17 | 100 | 99.88 | 100 | 100 | BXO1 | 99.87 | 4.5 | DRAFT | 63.7 | Rice | GCA_003296415.1 |
| X.oryzae pv. oryzae JL30 | 100 | 99.88 | 100 | 100 | BXO1 | 99.87 | 4 | DRAFT | 63 | Rice | GCA_003296355.1 |
| X.oryzae pv. oryzae JL24 | 100 | 99.88 | 100 | 100 | BXO1 | 99.87 | 4 | DRAFT | 62.9 | Rice | GCA_003296395.1 |
| X.oryzae pv. oryzae DBX0018 | 100 | 99.88 | 100 | 100 | BXO1 | 99.87 | 4.5 | DRAFT | 63.7 | Rice | GCA_003297945.1 |
| X.oryzae pv. oryzae LN18 | 100 | 99.88 | 100 | 100 | BXO1 | 99.87 | 5 | COMPLETE | 63.7 | Rice | GCA_009664105.1 |
| X.oryzae pv. oryzae HB84-21 | 100 | 99.88 | 100 | 100 | BXO1 | 99.87 | 3.9 | DRAFT | 62.7 | Rice | GCA_009664105.1 |
| X.oryzae pv. oryzae DBX0020 | 100 | 99.88 | 100 | 100 | BXO1 | 99.87 | 4.5 | DRAFT | 63.7 | Rice | GCA_003297895.1 |
| X.oryzae pv. oryzae DBX0016 | 100 | 99.88 | 100 | 100 | BXO1 | 99.87 | 4.5 | DRAFT | 63.7 | Rice | GCA_003295585.1 |
| X.oryzae pv. oryzae LN18 | 100 | 99.88 | 100 | 100 | BXO1 | 99.87 | 5 | DRAFT | 63.7 | Rice | GCA_009664105.1 |
| X.oryzae pv. oryzae JL4 | 100 | 99.88 | 100 | 100 | BXO1 | 99.87 | 4.5 | DRAFT | 63.7 | Rice | GCA_003296245.1 |
| X.oryzae pv. oryzae LN2 | 100 | 99.88 | 100 | 100 | BXO1 | 99.87 | 4.5 | DRAFT | 63.7 | Rice | GCA_003294875.1 |
| X.oryzae pv. oryzae JL1 | 100 | 99.88 | 100 | 100 | BXO1 | 99.87 | 4.5 | DRAFT | 63.7 | Rice | GCA_003296565.1 |
| X.oryzae pv. oryzae JL3 | 100 | 99.88 | 100 | 100 | BXO1 | 99.87 | 4.6 | DRAFT | 63.7 | Rice | GCA_003296305.1 |
| X.oryzae pv. oryzae LN3 | 100 | 99.88 | 100 | 100 | BXO1 | 99.87 | 4.5 | DRAFT | 63.7 | Rice | GCA_003295915.1 |
| X.oryzae pv. oryzae DBX0023 | 100 | 99.88 | 100 | 100 | BXO1 | 99.87 | 4.5 | DRAFT | 63.7 | Rice | GCA_003297905.1 |
| X.oryzae pv. oryzae DBX0024 | 100 | 99.88 | 100 | 100 | BXO1 | 99.87 | 4.5 | DRAFT | 63.7 | Rice | GCA_003297785.1 |
| X.oryzae pv. oryzae DBX004 | 100 | 99.88 | 100 | 100 | BXO1 | 99.87 | 4.5 | DRAFT | 63.7 | Rice | GCA_003297775.1 |
| X.oryzae pv. oryzae DBX0017 | 100 | 99.88 | 100 | 100 | BXO1 | 99.87 | 4.5 | DRAFT | 63.7 | Rice | GCA_003298065.1 |
| X.oryzae pv. oryzae YN03-14 | 100 | 99.88 | 100 | 100 | BXO1 | 99.87 | 4.6 | DRAFT | 63.7 | Rice | GCA_003299755.1 |
| X.oryzae pv. oryzae YN03-16 | 100 | 99.88 | 100 | 100 | BXO1 | 99.87 | 4.6 | DRAFT | 63.7 | Rice | GCA_003294335.1 |
| X.oryzae pv. oryzae YN20 | 100 | 99.88 | 100 | 100 | BXO1 | 99.87 | 4.6 | DRAFT | 63.7 | Rice | GCA_003294195.1 |
| X.oryzae pv. oryzae YN23 | 100 | 99.88 | 100 | 100 | BXO1 | 99.87 | 4.6 | DRAFT | 63.7 | Rice | GCA_003299265.1 |
| X.oryzae pv. oryzae YN03-13 | 100 | 99.88 | 100 | 100 | BXO1 | 99.87 | 4.6 | DRAFT | 63.7 | Rice | GCA_003299835.1 |
| X.oryzae pv. oryzae YN11 | 100 | 99.88 | 100 | 100 | BXO1 | 99.87 | 4.6 | DRAFT | 63.7 | Rice | GCA_002895775.1 |
| X.oryzae pv. oryzae YN18 | 100 | 99.88 | 100 | 100 | BXO1 | 99.87 | 4.6 | DRAFT | 63.7 | Rice | GCA_002895805.1 |
| X.oryzae pv. oryzae YN1 | 100 | 99.88 | 100 | 100 | BXO1 | 99.87 | 4.6 | DRAFT | 63.7 | Rice | GCA_002895745.1 |

| | | | | | | | | | | | |
|---|---|---|---|---|---|---|---|---|---|---|---|
| X.oryzae pv. oryzae YN44 | 100 | 99.88 | 100 | 100 | BXO1 | 99.87 | 4.6 | DRAFT | 63.7 | Rice | GCA_003299065.1 |
| X.oryzae pv. oryzae YN14 | 100 | 99.88 | 100 | 100 | BXO1 | 99.87 | 4.6 | DRAFT | 63.7 | Rice | GCA_003299395.1 |
| X.oryzae pv. oryzae YN8 | 100 | 99.88 | 100 | 100 | BXO1 | 99.87 | 4.6 | DRAFT | 63.7 | Rice | GCA_003294115.1 |
| X.oryzae pv. oryzae YN19 | 100 | 99.88 | 100 | 100 | BXO1 | 99.87 | 4.6 | DRAFT | 63.7 | Rice | GCA_003294205.1 |
| X.oryzae pv. oryzae YN5 | 100 | 99.88 | 100 | 100 | BXO1 | 99.87 | 4.6 | DRAFT | 63.7 | Rice | GCA_003299075.1 |
| X.oryzae pv. oryzae YN7 | 100 | 99.88 | 100 | 100 | BXO1 | 99.87 | 4.6 | DRAFT | 63.7 | Rice | GCA_002895765.1 |
| X.oryzae pv. oryzae YN6 | 100 | 99.88 | 100 | 100 | BXO1 | 99.87 | 4.6 | DRAFT | 63.7 | Rice | GCA_003294135.1 |
| X.oryzae pv. oryzae YN17 | 100 | 99.88 | 100 | 100 | BXO1 | 99.87 | 4.6 | DRAFT | 63.7 | Rice | GCA_003294215.1 |
| X.oryzae pv. oryzae YN40 | 100 | 99.88 | 100 | 100 | BXO1 | 99.87 | 4.6 | DRAFT | 63.7 | Rice | GCA_003294155.1 |
| X.oryzae pv. oryzae YN39 | 100 | 99.88 | 100 | 100 | BXO1 | 99.87 | 4.6 | DRAFT | 63.7 | Rice | GCA_003299145.1 |
| X.oryzae pv. oryzae ICMP3125 | 100 | 100 | 100 | 100 | BXO8 | 99.62 | 4.991 | COMPLETE | 63.7 | Rice | GCA_004136375.1 |
| X.oryzae pv. oryzae X035933 | 100 | 100 | 100 | 100 | BXO8 | 99.62 | 4.991 | COMPLETE | 63.7 | Rice | GCA_004136375.1 |
| X.oryzae pv. oryzae JS69 | 100 | 99.88 | 100 | 100 | BXO1 | 99.87 | 4.077 | DRAFT | 62.8 | Rice | GCA_003296295.1 |
| X.oryzae pv. oryzae PX079 | 100 | 99.88 | 100 | 100 | BXO1 | 99.87 | 5.027 | DRAFT | 63.7 | Rice | GCA_003382895.1 |
| X.oryzae pv. oryzae PXO124 | 100 | 99.88 | 100 | 100 | BXO1 | 99.87 | 4.432 | DRAFT | 63.3 | Rice | GCA_003300265.1 |
| X.oryzae pv. oryzae PXO99A | 100 | 99.88 | 100 | 100 | BXO1 | 99.87 | 5.239 | COMPLETE | 63.6 | Rice | GCA_000019585.2 |
| X.oryzae pv. oryzae DXO-015 | 100 | 99.88 | 100 | 100 | BXO1 | 99.87 | 4.275 | DRAFT | 63.9 | Rice | GCA_001927685.1 |
| X.oryzae pv. oryzae IXO411 | 100 | 99.88 | 100 | 100 | BXO1 | 99.87 | 4.303 | DRAFT | 63.9 | Rice | GCA_001928755.1 |
| X.oryzae pv. oryzae DXO-200 | 100 | 99.88 | 100 | 100 | BXO1 | 99.87 | 4.248 | DRAFT | 64 | Rice | GCA_001928445.1 |
| X.oryzae pv. oryzae IXO278 | 100 | 99.88 | 100 | 100 | BXO1 | 99.87 | 4.31 | DRAFT | 63.9 | Rice | GCA_001928185.1 |
| X.oryzae pv. oryzae IXO675 | 100 | 99.88 | 100 | 100 | BXO1 | 99.87 | 4.296 | DRAFT | 63.9 | Rice | GCA_001929075.1 |
| X.oryzae pv. oryzae DXO-242 | 100 | 99.88 | 100 | 100 | BXO1 | 99.87 | 4.276 | DRAFT | 64 | Rice | GCA_001927925.1 |
| X.oryzae pv. oryzae DXO-246 | 100 | 99.88 | 100 | 100 | BXO1 | 99.87 | 4.284 | DRAFT | 63.9 | Rice | GCA_001927945.1 |
| X.oryzae pv. oryzae YN04-5 | 100 | 99.88 | 100 | 100 | BXO1 | 99.87 | 4.551 | DRAFT | 63.8 | Rice | GCA_003299635.1 |
| X.oryzae pv. oryzae IXO884 | 100 | 99.88 | 100 | 100 | BXO1 | 99.87 | 4.325 | DRAFT | 63.9 | Rice | GCA_001929225.1 |
| X.oryzae pv. oryzae NX0260 | 100 | 100 | 100 | 100 | BXO8 | 99.62 | 5.05 | COMPLETE | 63.7 | Rice | GCA_004355825.1 |
| X.oryzae pv. oryzae BXO8 | 100 | 100 | 100 | 100 | BXO8 | 99.62 | 4.316 | DRAFT | 63.9 | Rice | GCA_001927415.1 |
| X.oryzae pv. oryzae CIAT | 100 | 98.66 | 100 | 99.33 | BXO8 | 99.62 | 5.105 | COMPLETE | 63.669 | Rice | GCA_004355865.1 |
| X.oryzae pv. oryzae IXO651 | 100 | 98.66 | 100 | 99.33 | BXO8 | 99.62 | 4.344 | DRAFT | 63.9 | Rice | GCA_001929065.1 |
| X.oryzae pv. oryzae IXO1221 | 100 | 98.66 | 100 | 99.33 | BXO8 | 99.62 | 4.343 | DRAFT | 63.9 | Rice | GCA_001929255.1 |
| X.oryzae pv. oryzae IXO685 | 100 | 98.66 | 100 | 99.33 | BXO8 | 99.62 | 4.372 | DRAFT | 63.8 | Rice | GCA_001929085.1 |
| X.oryzae pv. oryzae DXO-174 | 100 | 99.88 | 100 | 100 | BXO1 | 99.87 | 4.298 | DRAFT | 63.8 | Rice | GCA_001927855.1 |
| X.oryzae pv. oryzae DXO-233 | 100 | 99.88 | 100 | 100 | BXO1 | 99.87 | 4.381 | DRAFT | 63.9 | Rice | GCA_001928525.1 |
| X.oryzae pv. oryzae IXO1088 | 100 | 99.88 | 100 | 100 | BXO1 | 99.87 | 5.093 | DRAFT | 63.7 | Rice | GCA_001929235.2 |
| X.oryzae pv. oryzae IXO1104 | 100 | 99.88 | 100 | 100 | BXO1 | 99.87 | 4.394 | DRAFT | 63.8 | Rice | GCA_001929245.1 |
| X.oryzae pv. oryzae IXO725 | 100 | 99.88 | 100 | 100 | BXO1 | 99.87 | 4.379 | DRAFT | 63.8 | Rice | GCA_001929145.1 |
| X.oryzae pv. oryzae IXO89 | 100 | 99.88 | 100 | 100 | BXO1 | 99.87 | 4.297 | DRAFT | 63.9 | Rice | GCA_001928015.1 |
| X.oryzae pv. oryzae BXO6 | 100 | 99.88 | 100 | 100 | BXO1 | 99.87 | 4.274 | DRAFT | 63.9 | Rice | GCA_001927315.1 |
| X.oryzae pv. oryzae ITCCBB0002 | 100 | 99.88 | 100 | 100 | BXO1 | 99.87 | 4.732 | COMPLETE | 63.8 | Rice | GCA_009707765.1 |
| X.oryzae pv. oryzae IXO621 | 100 | 98.4 | 100 | 99.33 | BXO8 | 99.62 | 4.359 | DRAFT | 63.9 | Rice | GCA_001928935.1 |
| X.oryzae pv. oryzae IXO644 | 100 | 98.4 | 100 | 99.33 | BXO8 | 99.62 | 4.377 | DRAFT | 63.8 | Rice | GCA_001929015.1 |
| X.oryzae pv. oryzae IXO645 | 100 | 98.4 | 100 | 99.33 | BXO8 | 99.62 | 4.406 | DRAFT | 63.8 | Rice | GCA_001929005.1 |
| X.oryzae pv. oryzae IXO620 | 100 | 98.4 | 100 | 99.33 | BXO8 | 99.62 | 4.359 | DRAFT | 63.9 | Rice | GCA_001928915.1 |
| X.oryzae pv. oryzae IXO704 | 100 | 99.88 | 100 | 100 | BXO1 | 99.87 | 5.02 | DRAFT | 63.667 | Rice | GCA_001929095.2 |
| X.oryzae pv. oryzae IXO842 | 100 | 99.88 | 100 | 100 | BXO1 | 99.87 | 4.349 | DRAFT | 63.8 | Rice | GCA_001929165.1 |
| X.oryzae pv. oryzae IXO222 | 100 | 99.88 | 100 | 100 | BXO1 | 99.87 | 4.369 | DRAFT | 63.8 | Rice | GCA_001928685.1 |
| X.oryzae pv. oryzae IXO792 | 100 | 99.88 | 100 | 100 | BXO1 | 99.87 | 4.369 | DRAFT | 63.8 | Rice | GCA_001929155.1 |
| X.oryzae pv. oryzae IXO493 | 100 | 99.88 | 100 | 100 | BXO1 | 99.87 | 4.381 | DRAFT | 63.9 | Rice | GCA_001928825.1 |
| X.oryzae pv. oryzae IXO630 | 100 | 99.88 | 100 | 100 | BXO1 | 99.87 | 4.408 | DRAFT | 63.8 | Rice | GCA_001928985.1 |
| X.oryzae pv. oryzae IXO365 | 100 | 99.88 | 100 | 100 | BXO1 | 99.87 | 4.39 | DRAFT | 63.8 | Rice | GCA_001928695.1 |
| X.oryzae pv. oryzae IXO141 | 100 | 99.88 | 100 | 100 | BXO1 | 99.87 | 4.346 | DRAFT | 63.9 | Rice | GCA_001928675.1 |
| X.oryzae pv. oryzae IXO390 | 100 | 98.4 | 100 | 99.33 | BXO8 | 99.62 | 4.357 | DRAFT | 63.9 | Rice | GCA_001928765.1 |
| X.oryzae pv. oryzae IXO134 | 100 | 99.88 | 100 | 100 | BXO1 | 99.87 | 4.396 | DRAFT | 63.8 | Rice | GCA_001928095.1 |
| X.oryzae pv. oryzae BXO447 | 100 | 99.88 | 100 | 100 | BXO1 | 99.87 | 4.277 | DRAFT | 63.9 | Rice | GCA_001927545.1 |
| X.oryzae pv. oryzae BXO557 | 100 | 99.88 | 100 | 100 | BXO1 | 99.87 | 4.251 | DRAFT | 63.9 | Rice | GCA_001927625.1 |
| X.oryzae pv. oryzae YN24 | 100 | 98.4 | 100 | 99.33 | BXO8 | 99.62 | 4.476 | COMPLETE | 63.8 | Rice | GCA_003932075.1 |
| X.oryzae pv. oryzae IXO92 | 100 | 99.88 | 100 | 100 | BXO1 | 99.87 | 4.311 | DRAFT | 63.9 | Rice | GCA_001928615.1 |
| X.oryzae pv. oryzae BXO416 | 100 | 99.88 | 100 | 100 | BXO1 | 99.87 | 4.237 | DRAFT | 63.9 | Rice | GCA_001927495.1 |
| X.oryzae pv. oryzae BXO439 | 100 | 99.88 | 100 | 100 | BXO1 | 99.87 | 4.241 | DRAFT | 63.9 | Rice | GCA_001927305.1 |
| X.oryzae pv. oryzae DXO-170 | 100 | 99.88 | 100 | 100 | BXO1 | 99.87 | 4.328 | DRAFT | 63.9 | Rice | GCA_001928415.1 |
| X.oryzae pv. oryzae BXO34 | 100 | 99.88 | 100 | 100 | BXO1 | 99.87 | 4.287 | DRAFT | 63.9 | Rice | GCA_001927475.1 |
| X.oryzae pv. oryzae DXO-052 | 100 | 99.88 | 100 | 100 | BXO1 | 99.87 | 4.238 | DRAFT | 63.9 | Rice | GCA_001928325.1 |
| X.oryzae pv. oryzae DXO-089 | 100 | 99.88 | 100 | 100 | BXO1 | 99.87 | 4.293 | DRAFT | 63.9 | Rice | GCA_001927765.1 |

| | | | | | | | | | | | |
|---|---|---|---|---|---|---|---|---|---|---|---|
| X.oryzae pv. oryzae IXO220 | 100 | 99.88 | 100 | 100 | BXO1 | 99.87 | 4.299 | DRAFT | 63.9 | Rice | GCA_001928165.1 |
| X.oryzae pv. oryzae IXO627 | 100 | 99.88 | 100 | 100 | BXO1 | 99.87 | 4.332 | DRAFT | 63.9 | Rice | GCA_001928945.1 |
| X.oryzae pv. oryzae IXO189 | 100 | 99.88 | 100 | 100 | BXO1 | 99.87 | 4.281 | DRAFT | 63.9 | Rice | GCA_001928665.1 |
| X.oryzae pv. oryzae IXO639 | 100 | 99.88 | 100 | 100 | BXO1 | 99.87 | 4.275 | DRAFT | 63.9 | Rice | GCA_001928995.1 |
| X.oryzae pv. oryzae IXO98 | 100 | 99.88 | 100 | 100 | BXO1 | 99.87 | 4.256 | DRAFT | 63.9 | Rice | GCA_001928035.1 |
| X.oryzae pv. oryzae BXO33 | 100 | 99.88 | 100 | 100 | BXO1 | 99.87 | 4.274 | DRAFT | 63.9 | Rice | GCA_001927465.1 |
| X.oryzae pv. oryzae IXO151 | 100 | 99.88 | 100 | 100 | BXO1 | 99.87 | 4.2 | DRAFT | 63.9 | Rice | GCA_001928105.1 |
| X.oryzae pv. oryzae DXO-129 | 100 | 99.88 | 100 | 100 | BXO1 | 99.87 | 4.254 | DRAFT | 63.9 | Rice | GCA_001928365.1 |
| X.oryzae pv. oryzae DXO-165 | 100 | 99.88 | 100 | 100 | BXO1 | 99.87 | 4.304 | DRAFT | 63.9 | Rice | GCA_001927845.1 |
| X.oryzae pv. oryzae BXO-25 | 100 | 99.88 | 100 | 100 | BXO1 | 99.87 | 4.262 | DRAFT | 63.9 | Rice | GCA_001927395.1 |
| X.oryzae pv. oryzae DXO-027 | 100 | 99.88 | 100 | 100 | BXO1 | 99.87 | 4.224 | DRAFT | 63.9 | Rice | GCA_001927705.1 |
| X.oryzae pv. oryzae DXO-27 | 100 | 99.88 | 100 | 100 | BXO1 | 99.87 | 4.224 | DRAFT | 63.9 | Rice | GCA_001927705.1 |
| X.oryzae pv. oryzae BXO407 | 100 | 99.88 | 100 | 100 | BXO1 | 99.87 | 4.371 | DRAFT | 63.8 | Rice | GCA_001927335.1 |
| X.oryzae pv. oryzae BXO432 | 100 | 99.88 | 100 | 100 | BXO1 | 99.87 | 4.295 | DRAFT | 63.9 | Rice | GCA_001927505.1 |
| X.oryzae pv. oryzae IXO99 | 100 | 99.88 | 100 | 100 | BXO1 | 99.87 | 4.275 | DRAFT | 63.9 | Rice | GCA_001928085.1 |
| X.oryzae pv. oryzae BXO512 | 100 | 99.88 | 100 | 100 | BXO1 | 99.87 | 4.262 | DRAFT | 63.9 | Rice | GCA_001928215.2 |
| X.oryzae pv. oryzae DXO-044 | 100 | 99.88 | 100 | 100 | BXO1 | 99.87 | 4.251 | DRAFT | 63.9 | Rice | GCA_001927695.1 |
| X.oryzae pv. oryzae DXO-397 | 100 | 100 | 100 | 100 | BXO1 | 99.87 | 4.263 | DRAFT | 63.9 | Rice | GCA_001927955.1 |
| X.oryzae pv. oryzae SK2-3 | 100 | 99.88 | 100 | 100 | BXO1 | 99.87 | 4.934 | COMPLETE | 63.7 | Rice | GCA_003428965.1 |
| X.oryzae pv. oryzae IXO221 | 100 | 99.88 | 100 | 100 | BXO1 | 99.87 | 4.252 | DRAFT | 63.9 | Rice | GCA_001928175.1 |
| X.oryzae pv. oryzae IXO812 | 100 | 99.88 | 100 | 100 | BXO1 | 99.87 | 4.326 | DRAFT | 63.9 | Rice | GCA_001929185.1 |
| X.oryzae pv. oryzae DXO-050 | 100 | 99.88 | 100 | 100 | BXO1 | 99.87 | 4.355 | DRAFT | 63.9 | Rice | GCA_001927725.1 |
| X.oryzae pv. oryzae DXO-50 | 100 | 99.88 | 100 | 100 | BXO1 | 99.87 | 4.355 | DRAFT | 63.9 | Rice | GCA_001927725.1 |
| X.oryzae pv. oryzae DXO-122 | 100 | 100 | 100 | 100 | BXO1 | 99.87 | 4.263 | DRAFT | 64 | Rice | GCA_001927775.1 |
| X.oryzae pv. oryzae DXO-133 | 100 | 99.88 | 100 | 100 | BXO1 | 99.87 | 4.282 | DRAFT | 63.9 | Rice | GCA_001928775.1 |
| X.oryzae pv. oryzae IXO414 | 100 | 100 | 100 | 100 | BXO1 | 99.87 | 4.242 | DRAFT | 63.9 | Rice | GCA_001927795.1 |
| X.oryzae pv. oryzae BXO589 | 100 | 100 | 100 | 100 | BXO1 | 99.87 | 4.255 | DRAFT | 63.9 | Rice | GCA_001927605.1 |
| X.oryzae pv. oryzae BXO554 | 100 | 99.88 | 100 | 100 | BXO1 | 99.87 | 4.29 | DRAFT | 63.9 | Rice | GCA_001927385.1 |
| X.oryzae pv. oryzae BXO590 | 100 | 100 | 100 | 100 | BXO1 | 99.87 | 4.218 | DRAFT | 63.9 | Rice | GCA_001928265.1 |
| X.oryzae pv. oryzae DXO-116 | 100 | 100 | 100 | 100 | BXO1 | 99.87 | 4.2 | DRAFT | 63.9 | Rice | GCA_001928345.1 |
| X.oryzae pv. oryzae BXO1 | 100 | 99.88 | 100 | 100 | BXO1 | 99.87 | 5.084 | COMPLETE | 63.632 | Rice | GCA_001927405.2 |
| X.oryzae pv. oryzae BXO471 | 100 | 100 | 100 | 100 | BXO1 | 99.87 | 4.242 | DRAFT | 63.9 | Rice | GCA_001927615.1 |
| X.oryzae pv. oryzae DXO-091 | 100 | 100 | 100 | 100 | BXO1 | 99.87 | 4.29 | DRAFT | 63.9 | Rice | GCA_001928355.1 |
| X.oryzae pv. oryzae IXO367 | 100 | 100 | 100 | 100 | BXO1 | 99.87 | 4.254 | DRAFT | 63.9 | Rice | GCA_001928745.1 |
| X.oryzae pv. oryzae DXO-150 | 100 | 100 | 100 | 100 | BXO1 | 99.87 | 4.236 | DRAFT | 64 | Rice | GCA_001928405.1 |
| X.oryzae pv. oryzae DXO-248 | 100 | 100 | 100 | 100 | BXO1 | 99.87 | 4.228 | DRAFT | 63.9 | Rice | GCA_001928515.1 |
| X.oryzae pv. oryzae DXO-203 | 100 | 100 | 100 | 100 | BXO1 | 99.87 | 4.258 | DRAFT | 63.9 | Rice | GCA_001927865.1 |
| X.oryzae pv. oryzae DXO-369 | 100 | 100 | 100 | 100 | BXO1 | 99.87 | 4.262 | DRAFT | 63.9 | Rice | GCA_001928565.1 |
| X.oryzae pv. oryzae BXO2 | 100 | 99.88 | 100 | 100 | BXO1 | 99.87 | 4.239 | DRAFT | 63.9 | Rice | GCA_001927325.1 |
| X.oryzae pv. oryzae BXO558 | 100 | 99.88 | 100 | 100 | BXO1 | 99.87 | 4.259 | DRAFT | 63.9 | Rice | GCA_001928245.1 |
| X.oryzae pv. oryzae BXO582 | 100 | 100 | 100 | 100 | BXO1 | 99.87 | 4.272 | DRAFT | 63.9 | Rice | GCA_001928255.1 |
| X.oryzae pv. oryzae DXO-012 | 100 | 99.88 | 100 | 100 | BXO1 | 99.87 | 4.2 | DRAFT | 63.9 | Rice | GCA_001928285.1 |
| X.oryzae pv. oryzae BXO571 | 100 | 100 | 100 | 100 | BXO1 | 99.87 | 4.269 | DRAFT | 63.9 | Rice | GCA_001927575.1 |
| X.oryzae pv. oryzae BXO452 | 100 | 99.88 | 100 | 100 | BXO1 | 99.87 | 4.2 | DRAFT | 63.9 | Rice | GCA_001927575.1 |
| X.oryzae pv. oryzae BXO559 | 100 | 100 | 100 | 100 | BXO1 | 99.87 | 4.249 | DRAFT | 63.9 | Rice | GCA_001927635.1 |
| X.oryzae pv. oryzae IXO35 | 100 | 100 | 100 | 100 | BXO1 | 99.87 | 4.337 | DRAFT | 63.8 | Rice | GCA_001928585.1 |
| X.oryzae pv. oryzae DXO-133 | 100 | 100 | 100 | 100 | BXO1 | 99.87 | 4.282 | DRAFT | 63.9 | Rice | GCA_001927795.1 |
| X.oryzae pv. oryzae IXO74 | 100 | 99.88 | 100 | 100 | BXO1 | 99.87 | 4.314 | DRAFT | 63.9 | Rice | GCA_001928005.1 |
| X.oryzae pv. oryzae OS181 | 100 | 99.88 | 100 | 100 | BXO1 | 99.87 | 4.132 | DRAFT | 62.8 | Rice | GCA_003294675.1 |
| X.oryzae pv. oryzae G8 | 100 | 99.88 | 100 | 100 | BXO1 | 99.87 | 4.127 | DRAFT | 63.2 | Rice | GCA_003298095.1 |
| X.oryzae pv. oryzae YN38 | 100 | 99.88 | 100 | 100 | BXO1 | 99.87 | 4.22 | DRAFT | 63.2 | Rice | GCA_003299175.1 |
| X.oryzae pv. oryzae OS166 | 100 | 99.88 | 100 | 100 | BXO1 | 99.87 | 4.094 | DRAFT | 63.1 | Rice | GCA_003294695.1 |
| X.oryzae pv. oryzae YN16 | 100 | 99.88 | 100 | 100 | BXO1 | 99.87 | 4.544 | DRAFT | 63.8 | Rice | GCA_003299275.1 |
| X.oryzae pv. oryzae PXO61 | 100 | 99.88 | 100 | 100 | BXO1 | 99.87 | 4.999 | COMPLETE | 63.7 | Rice | GCA_004355885.3 |
| X.oryzae pv. oryzae PXO142 | 100 | 99.88 | 100 | 100 | BXO1 | 99.87 | 4.982 | COMPLETE | 63.7 | Rice | GCA_004136395.1 |
| X.oryzae pv. oryzae PXO513 | 100 | 99.88 | 100 | 100 | BXO1 | 99.87 | 4.916 | COMPLETE | 63.7 | Rice | GCA_004355705.1 |
| X.oryzae pv. oryzae PXO364 | 100 | 99.88 | 100 | 100 | BXO1 | 99.87 | 4.905 | COMPLETE | 63.7 | Rice | GCA_004355805.1 |
| X.oryzae pv. oryzae PXO421 | 100 | 99.88 | 100 | 100 | BXO1 | 99.87 | 4.91 | COMPLETE | 63.7 | Rice | GCA_004355725.1 |
| X.oryzae pv. oryzae PXO404 | 100 | 99.88 | 100 | 100 | BXO1 | 99.87 | 4.915 | COMPLETE | 63.7 | Rice | GCA_004355745.1 |
| X.oryzae pv. oryzae JS68 | 100 | 99.88 | 100 | 100 | BXO1 | 99.87 | 4.023 | DRAFT | 63 | Rice | GCA_003296035.1 |
| X.oryzae pv. oryzae 296 | 100 | 99.88 | 100 | 100 | BXO1 | 99.87 | 4.613 | DRAFT | 4.6129 | Rice | GCA_003298365.1 |
| X.oryzae pv. oryzae GD32 | 100 | 99.88 | 100 | 100 | BXO1 | 99.87 | 3.981 | DRAFT | 62.8 | Rice | GCA_003297535.1 |
| X.oryzae pv. oryzae XM-9 | 100 | 99.88 | 100 | 100 | BXO1 | 99.87 | 4.921 | COMPLETE | 63.7 | Rice | GCA_003522605.1 |

| | | | | | | | | | | | |
|---|---|---|---|---|---|---|---|---|---|---|---|
| X.oryzae pv. oryzae FuJ | 100 | 99.88 | 100 | 100 | BXO1 | 99.87 | 4.296 | DRAFT | 63.8 | Rice | GCA_002895665.1 |
| X.oryzae pv. oryzae HaN1 | 100 | 99.88 | 100 | 100 | BXO1 | 99.87 | 4.517 | DRAFT | 63.7 | Rice | GCA_003297035.1 |
| X.oryzae pv. oryzae OS-70 | 100 | 99.88 | 100 | 100 | BXO1 | 99.87 | 4.303 | DRAFT | 63.2 | Rice | GCA_003300385.1 |
| X.oryzae pv. oryzae GD38 | 100 | 99.88 | 100 | 100 | BXO1 | 99.87 | 4.272 | DRAFT | 63.2 | Rice | GCA_003297475.1 |
| X.oryzae pv. oryzae LFX325 | 100 | 99.88 | 100 | 100 | BXO1 | 99.87 | 3.979 | DRAFT | 62.8 | Rice | GCA_003294885.1 |
| X.oryzae pv. oryzae PXO563 | 100 | 99.88 | 100 | 100 | BXO1 | 99.87 | 4.936 | COMPLETE | 63.7 | Rice | GCA_001746715.1 |
| X.oryzae pv. oryzae HuN37 | 100 | 99.88 | 100 | 100 | BXO1 | 99.87 | 4.915 | COMPLETE | 63.7 | Rice | GCA_003382775.1 |
| X.oryzae pv. oryzae HuN33 | 100 | 99.88 | 100 | 100 | BXO1 | 99.87 | 4.369 | DRAFT | 63.5 | Rice | GCA_003296575.1 |
| X.oryzae pv. oryzae HuN38 | 100 | 99.88 | 100 | 100 | BXO1 | 99.87 | 4.204 | DRAFT | 63.2 | Rice | GCA_003296555.1 |
| X.oryzae pv. oryzae HuN32 | 100 | 99.88 | 100 | 100 | BXO1 | 99.87 | 4.402 | DRAFT | 63.5 | Rice | GCA_003296625.1 |
| X.oryzae pv. oryzae HuN36 | 100 | 99.88 | 100 | 100 | BXO1 | 99.87 | 4.403 | DRAFT | 63.5 | Rice | GCA_003296615.1 |
| X.oryzae pv. oryzae HuN39 | 100 | 99.88 | 100 | 100 | BXO1 | 99.87 | 4.212 | DRAFT | 63.2 | Rice | GCA_003295135.1 |
| X.oryzae pv. oryzae JP04 | 100 | 99.88 | 100 | 100 | BXO1 | 99.87 | 4.679 | DRAFT | 63.7 | Rice | GCA_003296105.1 |
| X.oryzae pv. oryzae JP03 | 100 | 99.88 | 100 | 100 | BXO1 | 99.87 | 4.603 | DRAFT | 63.7 | Rice | GCA_003296115.1 |
| X.oryzae pv. oryzae OS27 | 100 | 99.88 | 100 | 100 | BXO1 | 99.87 | 3.634 | DRAFT | 62.4 | Rice | GCA_003300505.1 |
| X.oryzae pv. oryzae HuN03-02 | 100 | 99.88 | 100 | 100 | BXO1 | 99.87 | 4.117 | DRAFT | 62.9 | Rice | GCA_003295215.1 |
| X.oryzae pv. oryzae AH1 | 100 | 99.88 | 100 | 100 | BXO1 | 99.87 | 4.635 | DRAFT | 63.8 | Rice | GCA_003298245.1 |
| X.oryzae pv. oryzae GX56 | 100 | 99.88 | 100 | 100 | BXO1 | 99.87 | 4.454 | DRAFT | 63.7 | Rice | GCA_003297085.1 |
| X.oryzae pv. oryzae DXO-181 | 100 | 99.88 | 100 | 100 | BXO1 | 99.87 | 4.276 | DRAFT | 63.9 | Rice | GCA_001928425.1 |
| X.oryzae pv. oryzae DXO-226 | 100 | 99.88 | 100 | 100 | BXO1 | 99.87 | 4.275 | DRAFT | 63.9 | Rice | GCA_001928495.1 |
| X.oryzae pv. oryzae GX6 | 100 | 99.88 | 100 | 100 | BXO1 | 99.87 | 4.463 | DRAFT | 63.8 | Rice | GCA_003295335.1 |
| X.oryzae pv. oryzae GX2 | 100 | 99.88 | 100 | 100 | BXO1 | 99.87 | 4.544 | DRAFT | 63.7 | Rice | GCA_003295375.1 |
| X.oryzae pv. oryzae GX1 | 100 | 99.88 | 100 | 100 | BXO1 | 99.87 | 4.57 | DRAFT | 63.7 | Rice | GCA_003297195.1 |
| X.oryzae pv. oryzae GX5 | 100 | 99.88 | 100 | 100 | BXO1 | 99.87 | 4.57 | DRAFT | 63.7 | Rice | GCA_003297095.1 |
| X.oryzae pv. oryzae GX3 | 100 | 99.88 | 100 | 100 | BXO1 | 99.87 | 4.57 | DRAFT | 63.7 | Rice | GCA_003297155.1 |
| X.oryzae pv. oryzae GX4 | 100 | 99.88 | 100 | 100 | BXO1 | 99.87 | 4.57 | DRAFT | 63.7 | Rice | GCA_003297145.1 |
| X.oryzae pv. oryzae LYG49 | 100 | 99.88 | 100 | 100 | BXO1 | 99.87 | 4.509 | DRAFT | 63.7 | Rice | GCA_003382265.1 |
| X.oryzae pv. oryzae FJ22 | 100 | 99.88 | 100 | 100 | BXO1 | 99.87 | 4.581 | DRAFT | 63.8 | Rice | GCA_003295525.1 |
| X.oryzae pv. oryzae YN08 | 100 | 99.88 | 100 | 100 | BXO1 | 99.87 | 4.157 | DRAFT | 63 | Rice | GCA_003299565.1 |
| X.oryzae pv. oryzae PXO71 | 100 | 99.88 | 100 | 100 | BXO1 | 99.87 | 3.633 | COMPLETE | 62.7 | Rice | GCA_003388465.1 |
| X.oryzae pv. oryzae DXO-331 | 100 | 99.88 | 100 | 100 | BXO1 | 99.87 | 4.319 | DRAFT | 63.9 | Rice | GCA_001927935.1 |
| X.oryzae pv. oryzae PXO602 | 100 | 99.88 | 100 | 100 | BXO1 | 99.87 | 4.952 | COMPLETE | 63.7 | Rice | GCA_001746735.1 |
| X.oryzae pv. oryzae PXO524 | 100 | 99.88 | 100 | 100 | BXO1 | 99.87 | 4.954 | COMPLETE | 63.7 | Rice | GCA_001746695.1 |
| X.oryzae pv. oryzae GD412 | 100 | 99.88 | 100 | 100 | BXO1 | 99.87 | 3.986 | DRAFT | 62.9 | Rice | GCA_003297265.1 |
| X.oryzae pv. oryzae GD401 | 100 | 99.88 | 100 | 100 | BXO1 | 99.87 | 3.986 | DRAFT | 62.9 | Rice | GCA_003295565.1 |
| X.oryzae pv. oryzae GD402 | 100 | 99.88 | 100 | 100 | BXO1 | 99.87 | 3.986 | DRAFT | 62.9 | Rice | GCA_003295545.1 |
| X.oryzae pv. oryzae GD408 | 100 | 99.88 | 100 | 100 | BXO1 | 99.87 | 3.986 | DRAFT | 62.9 | Rice | GCA_003297285.1 |
| X.oryzae pv. oryzae PXO282 | 100 | 99.88 | 100 | 100 | BXO1 | 99.87 | 4.962 | COMPLETE | 63.7 | Rice | GCA_001746675.1 |
| X.oryzae pv. oryzae GD404 | 100 | 99.88 | 100 | 100 | BXO1 | 99.87 | 4.677 | DRAFT | 63.7 | Rice | GCA_003297395.1 |
| X.oryzae pv. oryzae GD403 | 100 | 99.88 | 100 | 100 | BXO1 | 99.87 | 4.677 | DRAFT | 63.7 | Rice | GCA_003295515.1 |
| X.oryzae pv. oryzae GD416 | 100 | 99.88 | 100 | 100 | BXO1 | 99.87 | 4.677 | DRAFT | 63.7 | Rice | GCA_003295425.1 |
| X.oryzae pv. oryzae GD405 | 100 | 99.88 | 100 | 100 | BXO1 | 99.87 | 4.677 | DRAFT | 63.7 | Rice | GCA_003297375.1 |
| X.oryzae pv. oryzae GD414 | 100 | 99.88 | 100 | 100 | BXO1 | 99.87 | 4.677 | DRAFT | 63.7 | Rice | GCA_002895675.1 |
| X.oryzae pv. oryzae GD415 | 100 | 99.88 | 100 | 100 | BXO1 | 99.87 | 4.677 | DRAFT | 63.7 | Rice | GCA_003297245.1 |
| X.oryzae pv. oryzae GD413 | 100 | 99.88 | 100 | 100 | BXO1 | 99.87 | 4.677 | DRAFT | 63.7 | Rice | GCA_003295455.1 |
| X.oryzae pv. oryzae GD411 | 100 | 99.88 | 100 | 100 | BXO1 | 99.87 | 4.677 | DRAFT | 63.7 | Rice | GCA_003297275.1 |
| X.oryzae pv. oryzae GD417 | 100 | 99.88 | 100 | 100 | BXO1 | 99.87 | 4.677 | DRAFT | 63.7 | Rice | GCA_003297235.1 |
| X.oryzae pv. oryzae GD407 | 100 | 99.88 | 100 | 100 | BXO1 | 99.87 | 4.677 | DRAFT | 63.7 | Rice | GCA_003295485.1 |
| X.oryzae pv. oryzae GX23 | 100 | 99.88 | 100 | 100 | BXO1 | 99.87 | 4.135 | DRAFT | 63.2 | Rice | GCA_003297175.1 |
| X.oryzae pv. oryzae GX49 | 100 | 99.88 | 100 | 100 | BXO1 | 99.87 | 4.185 | DRAFT | 63.3 | Rice | GCA_003297135.1 |
| X.oryzae pv. oryzae GD33 | 100 | 99.88 | 100 | 100 | BXO1 | 99.87 | 4.589 | DRAFT | 63.7 | Rice | GCA_003297505.1 |
| X.oryzae pv. oryzae OS225 | 100 | 99.88 | 100 | 100 | BXO1 | 99.87 | 4.542 | DRAFT | 63.8 | Rice | GCA_003300515.1 |
| X.oryzae pv. oryzae OS221 | 100 | 100 | 100 | 100 | BXO1 | 99.87 | 4.542 | DRAFT | 63.8 | Rice | GCA_003300585.1 |
| X.oryzae pv. oryzae IX-280 | 100 | 99.88 | 100 | 100 | BXO1 | 99.87 | 5.007 | COMPLETE | 63.67 | Rice | GCA_003427055.1 |
| X.oryzae pv. oryzae XF89b | 100 | 99.88 | 100 | 100 | BXO1 | 99.87 | 4.967 | COMPLETE | 63.7 | Rice | GCA_002023005.1 |
| X.oryzae pv. oryzae JS72 | 100 | 99.88 | 100 | 100 | BXO1 | 99.87 | 3.949 | DRAFT | 62.8 | Rice | GCA_003294965.1 |
| X.oryzae pv. oryzae OS169 | 100 | 99.88 | 100 | 100 | BXO1 | 99.87 | 3.96 | DRAFT | 62.8 | Rice | GCA_003300815.1 |
| X.oryzae pv. oryzae DBX0013 | 100 | 99.88 | 100 | 100 | BXO1 | 99.87 | 4.485 | DRAFT | 63.7 | Rice | GCA_003298055.1 |
| X.oryzae pv. oryzae DBX0019 | 100 | 99.88 | 100 | 100 | BXO1 | 99.87 | 4.504 | DRAFT | 63.7 | Rice | GCA_003297915.1 |
| X.oryzae pv. oryzae IXO90 | 100 | 99.88 | 100 | 100 | BXO1 | 99.87 | 4.297 | DRAFT | 63.8 | Rice | GCA_001928605.1 |
| X.oryzae pv. oryzae IXO159 | 100 | 99.88 | 100 | 100 | BXO1 | 99.87 | 4.282 | DRAFT | 63.9 | Rice | GCA_001928135.1 |
| X.oryzae pv. oryzae IXO93 | 100 | 99.88 | 100 | 100 | BXO1 | 99.87 | 4.272 | DRAFT | 63.9 | Rice | GCA_001928025.1 |
| X.oryzae pv. oryzae IXO97 | 100 | 99.88 | 100 | 100 | BXO1 | 99.87 | 4.327 | DRAFT | 63.9 | Rice | GCA_001928625.1 |

| | | | | | | | | | | | |
|---|---|---|---|---|---|---|---|---|---|---|---|
| X.oryzae pv. oryzae DXO-206 | 100 | 99.88 | 100 | 100 | BXO1 | 99.87 | 4.378 | DRAFT | 63.9 | Rice | GCA_001928485.1 |
| X.oryzae pv. oryzae IXO603 | 100 | 99.88 | 100 | 100 | BXO1 | 99.87 | 4.279 | DRAFT | 63.9 | Rice | GCA_001928845.1 |
| X.oryzae pv. oryzae IXO608 | 100 | 99.88 | 100 | 100 | BXO1 | 99.87 | 4.276 | DRAFT | 63.9 | Rice | GCA_001928905.1 |
| X.oryzae pv. oryzae FJ17 | 100 | 99.88 | 100 | 100 | BXO1 | 99.87 | 4.588 | DRAFT | 63.7 | Rice | GCA_003297745.1 |
| X.oryzae pv. oryzae OS26 | 100 | 99.88 | 100 | 100 | BXO1 | 99.87 | 3.872 | DRAFT | 62.7 | Rice | GCA_003294635.1 |
| X.oryzae pv. oryzae HEN11 | 100 | 99.88 | 100 | 100 | BXO1 | 99.87 | 4.603 | DRAFT | 63.8 | Rice | GCA_003296805.1 |
| X.oryzae pv. oryzae HEN11 CTG0: | 100 | 99.88 | 100 | 100 | BXO1 | 99.87 | 4.603 | DRAFT | 63.8 | Rice | GCA_003296805.1 |
| X.oryzae pv. oryzae JP01 | 100 | 99.88 | 100 | 100 | BXO1 | 99.87 | 4.949 | COMPLETE | 63.7 | Rice | GCA_003294995.1 |
| X.oryzae pv. oryzae JP02 | 100 | 99.88 | 100 | 100 | BXO1 | 99.87 | 4.949 | DRAFT | 63.7 | Rice | GCA_003382855.1 |
| X.oryzae pv. oryzae MAFF311018 | 100 | 99.88 | 100 | 100 | BXO1 | 99.87 | 4.94 | COMPLETE | 63.7 | Rice | GCA_000010025.1 |
| X.oryzae pv. oryzae LYG46 | 100 | 99.88 | 100 | 100 | BXO1 | 99.87 | 4.533 | DRAFT | 63.8 | Rice | GCA_003295805.1 |
| X.oryzae pv. oryzae OS35 | 100 | 99.88 | 100 | 100 | BXO1 | 99.87 | 3.836 | DRAFT | 62.6 | Rice | GCA_003294605.1 |
| X.oryzae pv. oryzae OS54 | 100 | 99.88 | 100 | 100 | BXO1 | 99.87 | 3.876 | DRAFT | 62.7 | Rice | GCA_003294585.1 |
| X.oryzae pv. oryzae OS34 | 100 | 99.88 | 100 | 100 | BXO1 | 99.87 | 3.718 | DRAFT | 62.5 | Rice | GCA_003300465.1 |
| X.oryzae pv. oryzae OS44 | 100 | 99.88 | 100 | 100 | BXO1 | 99.87 | 3.862 | DRAFT | 62.6 | Rice | GCA_003294595.1 |
| X.oryzae pv. oryzae SX-85-73 | 100 | 99.88 | 100 | 100 | BXO1 | 99.87 | 4.596 | DRAFT | 63.8 | Rice | GCA_003294505.1 |
| X.oryzae pv. oryzae JX19 | 100 | 99.88 | 100 | 100 | BXO1 | 99.87 | 4.578 | DRAFT | 63.8 | Rice | GCA_003294955.1 |
| X.oryzae pv. oryzae HuN17 | 100 | 99.88 | 100 | 100 | BXO1 | 99.87 | 4.261 | DRAFT | 63.3 | Rice | GCA_003296675.1 |
| X.oryzae pv. oryzae HuN20 | 100 | 99.88 | 100 | 100 | BXO1 | 99.87 | 4.552 | DRAFT | 63.7 | Rice | GCA_003295185.1 |
| X.oryzae pv. oryzae HB84-17 | 100 | 99.88 | 100 | 100 | BXO1 | 99.87 | 3.965 | DRAFT | 62.8 | Rice | GCA_003295325.1 |
| X.oryzae pv. oryzae HEN03-02 | 100 | 99.88 | 100 | 100 | BXO1 | 99.87 | 3.942 | DRAFT | 62.8 | Rice | GCA_003296945.1 |
| X.oryzae pv. oryzae HeN03-03 | 100 | 99.88 | 100 | 100 | BXO1 | 99.87 | 3.942 | DRAFT | 62.8 | Rice | GCA_003296935.1 |
| X.oryzae pv. oryzae YN03-10 | 100 | 99.88 | 100 | 100 | BXO1 | 99.87 | 4.598 | DRAFT | 63.7 | Rice | GCA_003299865.1 |
| X.oryzae pv. oryzae HeN03-16 | 100 | 99.88 | 100 | 100 | BXO1 | 99.87 | 4.395 | DRAFT | 63.4 | Rice | GCA_003296845.1 |
| X.oryzae pv. oryzae HeN03-11 | 100 | 99.88 | 100 | 100 | BXO1 | 99.87 | 4.395 | DRAFT | 63.4 | Rice | GCA_003296865.1 |
| X.oryzae pv. oryzae HeN03-10 | 100 | 99.88 | 100 | 100 | BXO1 | 99.87 | 4.395 | DRAFT | 63.4 | Rice | GCA_003295275.1 |
| X.oryzae pv. oryzae HeN03-06 | 100 | 99.88 | 100 | 100 | BXO1 | 99.87 | 4.395 | DRAFT | 63.4 | Rice | GCA_003295315.1 |
| X.oryzae pv. oryzae HeN03-04 | 100 | 99.88 | 100 | 100 | BXO1 | 99.87 | 4.395 | DRAFT | 63.4 | Rice | GCA_003296925.1 |
| X.oryzae pv. oryzae HeN03-01 | 100 | 99.88 | 100 | 100 | BXO1 | 99.87 | 4.395 | DRAFT | 63.4 | Rice | GCA_003296915.1 |
| X.oryzae pv. oryzae KSI-3 | 100 | 99.88 | 100 | 100 | BXO1 | 99.87 | 4.326 | DRAFT | 63.3 | Rice | GCA_003295975.1 |
| X.oryzae pv. oryzae 296 | 100 | 99.88 | 100 | 100 | BXO1 | 99.87 | 4.613 | DRAFT | 4.6129 | Rice | GCA_003298365.1 |
| X.oryzae pv. oryzae FJ24 | 100 | 99.88 | 100 | 100 | BXO1 | 99.87 | 4.63 | DRAFT | 63.7 | Rice | GCA_003297625.1 |
| X.oryzae pv. oryzae LYG03-13 | 100 | 99.88 | 100 | 100 | BXO1 | 99.87 | 4.53 | DRAFT | 63.8 | Rice | GCA_003294745.1 |
| X.oryzae pv. oryzae LYG51 | 100 | 99.88 | 100 | 100 | BXO1 | 99.87 | 4.53 | DRAFT | 63.8 | Rice | GCA_003295775.1 |
| X.oryzae pv. oryzae LYG53 | 100 | 99.88 | 100 | 100 | BXO1 | 99.87 | 4.53 | DRAFT | 63.8 | Rice | GCA_003300845.1 |
| X.oryzae pv. oryzae LYG52 | 100 | 99.88 | 100 | 100 | BXO1 | 99.87 | 4.53 | DRAFT | 63.8 | Rice | GCA_003294715.1 |
| X.oryzae pv. oryzae K3 | 100 | 99.88 | 100 | 100 | BXO1 | 99.87 | 4.946 | COMPLETE | 63.7 | Rice | GCA_011604765.1 |
| X.oryzae pv. oryzae OS182 | 100 | 99.88 | 100 | 100 | BXO1 | 99.87 | 4.399 | DRAFT | 63.3 | Rice | GCA_003294665.1 |
| X.oryzae pv. oryzae OS82 | 100 | 99.88 | 100 | 100 | BXO1 | 99.87 | 4.077 | DRAFT | 62.9 | Rice | GCA_003300375.1 |
| X.oryzae pv. oryzae OS77 | 100 | 99.88 | 100 | 100 | BXO1 | 99.87 | 4.461 | DRAFT | 63.4 | Rice | GCA_003294575.1 |
| X.oryzae pv. oryzae HN-7 | 100 | 99.88 | 100 | 100 | BXO1 | 99.87 | 4.27 | DRAFT | 63.3 | Rice | GCA_003295235.1 |
| X.oryzae pv. oryzae KS2-2 | 100 | 99.88 | 100 | 100 | BXO1 | 99.87 | 4.014 | DRAFT | 62.9 | Rice | GCA_003294945.1 |
| X.oryzae pv. oryzae HB84-35 | 100 | 99.88 | 100 | 100 | BXO1 | 99.87 | 3.9 | DRAFT | 62.8 | Rice | GCA_003296685.1 |
| X.oryzae pv. oryzae HEN08 | 100 | 99.88 | 100 | 100 | BXO1 | 99.87 | 4.301 | DRAFT | 63.2 | Rice | GCA_003296855.1 |
| X.oryzae pv. oryzae JS65 | 100 | 99.88 | 100 | 100 | BXO1 | 99.87 | 4.534 | DRAFT | 63.7 | Rice | GCA_003296065.1 |
| X.oryzae pv. oryzae HEN12 | 100 | 99.88 | 100 | 100 | BXO1 | 99.87 | 4 | DRAFT | 62.6 | Rice | GCA_003296815.1 |
| X.oryzae pv. oryzae BJ84-31 | 100 | 99.88 | 100 | 100 | BXO1 | 99.87 | 4 | DRAFT | 62.6 | Rice | GCA_003298095.1 |
| X.oryzae pv. oryzae OS15 | 100 | 99.88 | 100 | 100 | BXO1 | 99.87 | 3.9 | DRAFT | 62.8 | Rice | GCA_003301335.1 |
| X.oryzae pv. oryzae HUN22 | 100 | 99.88 | 100 | 100 | BXO1 | 99.87 | 4.5 | DRAFT | 63.8 | Rice | GCA_003295175.1 |
| X.oryzae pv. oryzae OS48 | 100 | 99.88 | 100 | 100 | BXO1 | 99.87 | 3.9 | DRAFT | 62.7 | Rice | GCA_003301295.1 |
| X.oryzae pv. oryzae HUB14 | 100 | 99.88 | 100 | 100 | BXO1 | 99.87 | 4.5 | DRAFT | 63.8 | Rice | GCA_003296695.1 |
| X.oryzae pv. oryzae JNXO | 100 | 99.88 | 100 | 100 | BXO1 | 99.87 | 4.1 | DRAFT | 63.2 | Rice | GCA_003296135.1 |
| X.oryzae pv. oryzae AH3 | 100 | 99.88 | 100 | 100 | BXO1 | 99.87 | 3.8 | DRAFT | 62.9 | Rice | GCA_003298235.1 |
| X.oryzae pv. oryzae K3a | 100 | 99.88 | 100 | 100 | BXO1 | 99.87 | 4.8 | COMPLETE | 63.7 | Rice | GCA_011604785.1 |
| X.oryzae pv. oryzae ScHy-b | 100 | 99.88 | 100 | 100 | BXO1 | 99.87 | 4.5 | DRAFT | 63.7 | Rice | GCA_003294525.1 |
| X.oryzae pv. oryzae ScYc-b | 100 | 99.88 | 100 | 100 | BXO1 | 99.87 | 4.8 | COMPLETE | 63.7 | Rice | GCA_002895725.2 |
| X.oryzae pv. oryzae OS185 | 100 | 99.88 | 100 | 100 | BXO1 | 99.87 | 4.4 | DRAFT | 63.4 | Rice | GCA_003300715.1 |
| X.oryzae pv. oryzae JS158-2 | 100 | 99.88 | 100 | 100 | BXO1 | 99.87 | 4 | DRAFT | 63.1 | Rice | GCA_003296075.1 |
| X.oryzae pv. oryzae ZJ05 | 100 | 99.88 | 100 | 100 | BXO1 | 99.87 | 4.5 | DRAFT | 63.8 | Rice | GCA_003299055.1 |
| X.oryzae pv. oryzae HUB03-06 | 100 | 99.88 | 100 | 100 | BXO1 | 99.87 | 4 | DRAFT | 62.9 | Rice | GCA_003295225.1 |
| X.oryzae pv. oryzae ZJ16 | 100 | 99.88 | 100 | 100 | BXO1 | 99.87 | 4.2 | DRAFT | 63.2 | Rice | GCA_003294095.1 |
| X.oryzae pv. oryzae ZJ27 | 100 | 99.88 | 100 | 100 | BXO1 | 99.87 | 4.5 | DRAFT | 63.7 | Rice | GCA_003298965.1 |
| X.oryzae pv. oryzae ZJ26 | 100 | 99.88 | 100 | 100 | BXO1 | 99.87 | 4.2 | DRAFT | 63.3 | Rice | GCA_003298955.1 |

| | | | | | | | | | | | |
|---|---|---|---|---|---|---|---|---|---|---|---|
| X.oryzae pv. oryzae ZJ25 | 100 | 99.88 | 100 | 100 | BXO1 | 99.87 | 4.6 | DRAFT | 63.8 | Rice | GCA_003294085.1 |
| X.oryzae pv. oryzae AH8 | 100 | 99.88 | 100 | 100 | BXO1 | 99.87 | 4.6 | DRAFT | 63.8 | Rice | GCA_003295635.1 |
| X.oryzae pv. oryzae AH40 | 100 | 99.88 | 100 | 100 | BXO1 | 99.87 | 3.9 | DRAFT | 628 | Rice | GCA_003298195.1 |
| X.oryzae pv. oryzae AH11 | 100 | 99.88 | 100 | 100 | BXO1 | 99.87 | 4.6 | DRAFT | 63.8 | Rice | GCA_003298245.1 |
| X.oryzae pv. oryzae AH87 | 100 | 99.88 | 100 | 100 | BXO1 | 99.87 | 4.4 | DRAFT | 63.4 | Rice | GCA_003295635.1 |
| X.oryzae pv. oryzae AH39 | 100 | 99.88 | 100 | 100 | BXO1 | 99.87 | 4.1 | DRAFT | 62.9 | Rice | GCA_003298235.1 |
| X.oryzae pv. oryzae ZJ28 | 100 | 99.88 | 100 | 100 | BXO1 | 99.87 | 4.6 | DRAFT | 63.7 | Rice | GCA_003298825.1 |
| X.oryzae pv. oryzae XZ44 | 100 | 99.88 | 100 | 100 | BXO1 | 99.87 | 4.5 | DRAFT | 63.7 | Rice | GCA_003300125.1 |
| X.oryzae pv. oryzae XZ43 | 100 | 99.88 | 100 | 100 | BXO1 | 99.87 | 4.5 | DRAFT | 63.7 | Rice | GCA_003294465.1 |
| X.oryzae pv. oryzae XZ42 | 100 | 99.88 | 100 | 100 | BXO1 | 99.87 | 4.5 | DRAFT | 63.7 | Rice | GCA_003300215.1 |
| X.oryzae pv. oryzae XZ39 | 100 | 99.88 | 100 | 100 | BXO1 | 99.87 | 4.5 | DRAFT | 63.7 | Rice | GCA_003300225.1 |
| X.oryzae pv. oryzae XZ40 | 100 | 99.88 | 100 | 100 | BXO1 | 99.87 | 4.5 | DRAFT | 63.7 | Rice | GCA_003294495.1 |
| X.oryzae pv. oryzae LYG48 | 100 | 99.88 | 100 | 100 | BXO1 | 99.87 | 4.5 | DRAFT | 63.8 | Rice | GCA_003295795.1 |
| X.oryzae pv. oryzae YC36 | 100 | 99.88 | 100 | 100 | BXO1 | 99.87 | 4.5 | DRAFT | 63.8 | Rice | GCA_003300055.1 |
| X.oryzae pv. oryzae YC20 | 100 | 99.88 | 100 | 100 | BXO1 | 99.87 | 4.5 | DRAFT | 63.8 | Rice | GCA_003300075.1 |
| X.oryzae pv. oryzae YC26 | 100 | 99.88 | 100 | 100 | BXO1 | 99.87 | 4.4 | DRAFT | 63.7 | Rice | GCA_003294445.1 |
| X.oryzae pv. oryzae ZPY1 | 100 | 99.88 | 100 | 100 | BXO1 | 99.87 | 4.1 | DRAFT | 63.2 | Rice | GCA_003298815.1 |
| X.oryzae pv. oryzae AH4 | 100 | 99.88 | 100 | 100 | BXO1 | 99.87 | 4 | DRAFT | 62.8 | Rice | GCA_003298205.1 |
| X.oryzae pv. oryzae YC11 | 100 | 99.88 | 100 | 100 | BXO1 | 99.87 | 4.8 | COMPLETE | 63.7 | Rice | GCA_003382935.1 |
| X.oryzae pv. oryzae YC3 | 100 | 99.88 | 100 | 100 | BXO1 | 99.87 | 4.4 | DRAFT | 63.8 | Rice | GCA_003294425.1 |
| X.oryzae pv. oryzae LYG03-05 | 100 | 99.88 | 100 | 100 | BXO1 | 99.87 | 4.1 | DRAFT | 63 | Rice | GCA_003294765.1 |
| X.oryzae pv. oryzae LYG03-08 | 100 | 99.88 | 100 | 100 | BXO1 | 99.87 | 4.1 | DRAFT | 63 | Rice | GCA_003295855.1 |
| X.oryzae pv. oryzae LYG03-02 | 100 | 99.88 | 100 | 100 | BXO1 | 99.87 | 4.1 | DRAFT | 63 | Rice | GCA_003295855.1 |
| X.oryzae pv. oryzae LYG03-11 | 100 | 99.88 | 100 | 100 | BXO1 | 99.87 | 4.1 | DRAFT | 63 | Rice | GCA_003295845.1 |
| X.oryzae pv. oryzae 172 | 100 | 99.88 | 100 | 100 | BXO1 | 99.87 | 4.6 | DRAFT | 63.7 | Rice | GCA_003363805.1 |
| X.oryzae pv. oryzae HN14 | 100 | 99.88 | 100 | 100 | BXO1 | 99.87 | 4.2 | DRAFT | 63.2 | Rice | GCA_003296745.1 |
| X.oryzae pv. oryzae HUN14 | 100 | 99.88 | 100 | 100 | BXO1 | 99.87 | 4.1 | DRAFT | 63.2 | Rice | GCA_003296745.1 |
| X.oryzae pv. oryzae GX07 | 100 | 99.88 | 100 | 100 | BXO1 | 99.87 | 3.9 | DRAFT | 62.9 | Rice | GCA_003295435.1 |
| X.oryzae pv. oryzae GX09 | 100 | 99.88 | 100 | 100 | BXO1 | 99.87 | 4.1 | DRAFT | 63.2 | Rice | GCA_003295405.1 |
| X.oryzae pv. oryzae HLJ02 | 100 | 99.88 | 100 | 100 | BXO1 | 99.87 | 4.4 | DRAFT | 63.8 | Rice | GCA_003295245.1 |
| X.oryzae pv. oryzae LN85-47 | 100 | 99.88 | 100 | 100 | BXO1 | 99.87 | 3.9 | DRAFT | 62.8 | Rice | GCA_003295865.1 |
| X.oryzae pv. oryzae HLJ85-69 | 100 | 99.88 | 100 | 100 | BXO1 | 99.87 | 4.1 | DRAFT | 63.2 | Rice | GCA_003296775.1 |
| X.oryzae pv. oryzae LYG03-04 | 100 | 99.88 | 100 | 100 | BXO1 | 99.87 | 4.1 | DRAFT | 63 | Rice | GCA_003294795.1 |
| X.oryzae pv. oryzae LYG03-07 | 100 | 99.88 | 100 | 100 | BXO1 | 99.87 | 4.1 | DRAFT | 63 | Rice | GCA_003295825.1 |
| X.oryzae pv. oryzae LN01 | 100 | 99.88 | 100 | 100 | BXO1 | 99.87 | 4.51 | DRAFT | 63.8 | Rice | GCA_003295965.1 |
| X.oryzae pv. oryzae LN02 | 100 | 99.88 | 100 | 100 | BXO1 | 99.87 | 4.4 | DRAFT | 63.8 | Rice | GCA_003295955.1 |
| X.oryzae pv. oryzae LN85-59 | 100 | 99.88 | 100 | 100 | BXO1 | 99.87 | 4.4 | DRAFT | 63.8 | Rice | GCA_003294835.1 |
| X.oryzae pv. oryzae JL22 | 100 | 99.88 | 100 | 100 | BXO1 | 99.87 | 4.8 | DRAFT | 63.8 | Rice | GCA_003296405.1 |
| X.oryzae pv. oryzae JL6 | 100 | 99.88 | 100 | 100 | BXO1 | 99.87 | 4.8 | DRAFT | 63.8 | Rice | GCA_003295045.1 |
| X.oryzae pv. oryzae JL33 | 100 | 99.88 | 100 | 100 | BXO1 | 99.87 | 4.8 | COMPLETE | 63.8 | Rice | GCA_003382835.1 |
| X.oryzae pv. oryzae JL31 | 100 | 99.88 | 100 | 100 | BXO1 | 99.87 | 4.9 | DRAFT | 63.8 | Rice | GCA_003296335.1 |
| X.oryzae pv. oryzae JL25 | 100 | 99.88 | 100 | 100 | BXO1 | 99.87 | 4.8 | COMPLETE | 63.7 | Rice | GCA_003382795.1 |
| X.oryzae pv. oryzae JL23 | 100 | 99.88 | 100 | 100 | BXO1 | 99.87 | 4.9 | DRAFT | 63.7 | Rice | GCA_003295065.1 |
| X.oryzae pv. oryzae JL21 | 100 | 99.88 | 100 | 100 | BXO1 | 99.87 | 4.9 | DRAFT | 63.7 | Rice | GCA_003296255.1 |
| X.oryzae pv. oryzae JL 1 2003 | 100 | 99.88 | 100 | 100 | BXO1 | 99.87 | 4.9 | DRAFT | 63.7 | Rice | GCA_003296505.1 |
| X.oryzae pv. oryzae JL19 | 100 | 99.88 | 100 | 100 | BXO1 | 99.87 | 4.9 | DRAFT | 63.7 | Rice | GCA_003295075.1 |
| X.oryzae pv. oryzae JL13 | 100 | 99.88 | 100 | 100 | BXO1 | 99.87 | 4.9 | DRAFT | 63.7 | Rice | GCA_003296495.1 |
| X.oryzae pv. oryzae JL28 | 100 | 99.88 | 100 | 100 | BXO1 | 99.87 | 4.7 | COMPLETE | 63.7 | Rice | GCA_003382815.1 |
| X.oryzae pv. oryzae JL11 | 100 | 99.88 | 100 | 100 | BXO1 | 99.87 | 4.9 | DRAFT | 63.7 | Rice | GCA_003296515.1 |
| X.oryzae pv. oryzae JL12 | 100 | 99.88 | 100 | 100 | BXO1 | 99.87 | 4.9 | DRAFT | 63.7 | Rice | GCA_003296465.1 |
| X.oryzae pv. oryzae JL15 | 100 | 99.88 | 100 | 100 | BXO1 | 99.87 | 4.9 | DRAFT | 63.7 | Rice | GCA_003296455.1 |
| X.oryzae pv. oryzae JL16 | 100 | 99.88 | 100 | 100 | BXO1 | 99.87 | 4.9 | DRAFT | 63.7 | Rice | GCA_003295085.1 |
| X.oryzae pv. oryzae JL29 | 100 | 99.88 | 100 | 100 | BXO1 | 99.87 | 4.9 | DRAFT | 63.7 | Rice | GCA_003295035.1 |
| X. axonopodis DSM 3585 | 100 | 94.64 | 100 | 94 | BXO8 | | 4.48 | Contig | 64.5 | Axonopı | GCA_001304695.1 |
| X. axonopodis Xa85 | 100 | 94.64 | 100 | 94 | BXO8 | | 4.258 | Scaffold | 64.5 | Axonopı | GCA_003111925.1 |
| X. axonopodis pv. vasculorum NC | 100 | 94.64 | 100 | 94 | BXO8 | | 4.8 | Scaffold | 64 | Axonopı | GCA_000724905.2 |
| X. axonopodis pv. vasculorum CFI | 100 | 94.64 | 100 | 94 | BXO8 | | 4.692 | Contig | 64.2 | Axonopı | GCA_002939725.1 |
| X. axonopodis pv. vasculorum NC | 100 | 94.64 | 100 | 94 | BXO8 | | 4.887 | Complete | 64.089 | Axonopı | GCA_013177355.1 |
| X. vasicola CFBP2543 | 100 | 99.75 | 100 | 92.2 | Xvv | 99.96 | 4.922 | Contig | 63.3 | Ensete v | GCA_002939925.1 |
| X. vasicola NCPPB 2417 | 100 | 99.75 | 100 | 92.2 | Xvv | 99.96 | 4.924 | Contig | 63.3 | Ensete v | GCA_000772705.2 |
| X. vasicola Mex-1 | 100 | 99.75 | 100 | 92.2 | Xvv | 99.96 | 4.987 | Contig | 63.3 | Ensete v | GCA_007846165.1 |
| X. vasicola CO-5 | 100 | 100 | 100 | 100 | Xvv | 99.96 | 4.957 | Contig | 63.2 | Ensete v | GCA_007846415.1 |
| X. vasicola NE744 | 100 | 100 | 100 | 100 | Xvv | 99.96 | 4.957 | Contig | 63.2 | Ensete v | GCA_007846185.1 |

| | | | | | | | | | | |
|---|---|---|---|---|---|---|---|---|---|---|
| X. vasicola ZCP611 | 84 | 67.77 | 94 | 69.24 | BXO8 | 94.3 | 5.018 | Contig | 63.3 | sugarcar GCA_007846255.1 |
| X. vasicola BCC250 | 84 | 67.77 | 94 | 69.24 | BXO8 | 94.3 | 4.713 | Contig | 63.5 | sugarcar GCA_003312845.2 |
| X. vasicola BCC274 | 84 | 67.77 | 94 | 69.24 | BXO8 | 94.3 | 4.754 | Contig | 63.5 | sugarcar GCA_003312695.2 |
| X. vasicola NE-3 | 100 | 100 | 100 | 100 | Xvv | 99.96 | 4.945 | Scaffold | 63.2 | Ensete v GCA_007846405.1 |
| X. vasicola NE-5 | 100 | 100 | 100 | 100 | Xvv | 99.96 | 4.939 | Scaffold | 63.2 | Ensete v GCA_007846465.1 |
| X. vasicola Arg-3B | 100 | 100 | 100 | 100 | Xvv | 99.96 | 4.937 | Scaffold | 63.2 | Ensete v GCA_007846145.1 |
| X. vasicola 4W4 | 100 | 100 | 100 | 100 | Xvv | 99.96 | 4.751 | Contig | 63.5 | Ensete v GCA_003312795.4 |
| X. vasicola BCC210 | 100 | 100 | 100 | 100 | Xvv | 99.96 | 4.752 | Contig | 63.5 | Ensete v GCA_003312785.2 |
| X. vasicola BCC246 | 84 | 67.77 | 94 | 69.24 | BXO8 | 94.3 | 4.755 | Contig | 63.5 | sugarcar GCA_003312775.3 |
| X. vasicola BCC278 | 84 | 67.77 | 94 | 69.24 | BXO8 | 94.3 | 4.715 | Contig | 63.5 | sugarcar GCA_003312685.2 |
| X. vasicola BCC280 | 84 | 67.77 | 94 | 69.24 | BXO8 | 94.3 | 4.716 | Contig | 63.5 | sugarcar GCA_003312615.2 |
| X. vasicola BCC248 | 84 | 67.77 | 94 | 69.24 | BXO8 | 94.3 | 4.752 | Contig | 63.5 | sugarcar GCA_003312715.2 |
| X. vasicola BCC265 | 84 | 67.77 | 94 | 69.24 | BXO8 | 94.3 | 4.752 | Contig | 63.5 | sugarcar GCA_003312705.2 |
| X. vasicola BCC281 | 84 | 67.77 | 94 | 69.24 | BXO8 | 94.3 | 4.718 | Contig | 63.5 | sugarcar GCA_003312625.2 |
| X. vasicola R5P | 84 | 67.77 | 94 | 69.24 | BXO8 | 94.3 | 4.712 | Contig | 63.5 | sugarcar GCA_003312605.3 |
| X. vasicola BCC247 | 84 | 67.77 | 94 | 69.24 | BXO8 | 94.3 | 4.756 | Contig | 63.5 | sugarcar GCA_003312765.2 |
| X. vasicola DAR-82621 | 100 | 99.75 | 100 | 99.22 | Xvv | 99.96 | 4.924 | Contig | 63.3 | Ensete v GCA_007846515.1 |
| X. vasicola DAR-41380 | 100 | 99.75 | 100 | 99.22 | Xvv | 99.96 | 4.989 | Contig | 63.3 | Ensete v GCA_007846295.1 |
| X. vasicola Xvh-L | 100 | 99.75 | 100 | 99.22 | Xvv | 99.96 | 5.273 | Scaffold | 62.9 | Ensete v GCA_007846385.1 |
| X. vasicola DAR-82637 | 100 | 99.75 | 100 | 99.22 | Xvv | 99.96 | 4.979 | Contig | 63.3 | Ensete v GCA_007846155.1 |
| X. vasicola DAR-82872 | 100 | 99.75 | 100 | 99.22 | Xvv | 99.96 | 4.886 | Contig | 63.4 | Ensete v GCA_007846395.1 |
| X. vasicola BRIP-39150 | 100 | 99.75 | 100 | 99.22 | Xvv | 99.96 | 4.953 | Contig | 63.3 | Ensete v GCA_007846235.1 |
| X. vasicola DAR-82728 | 100 | 99.75 | 100 | 99.22 | Xvv | 99.96 | 4.883 | Contig | 63.4 | Ensete v GCA_007846245.1 |
| X. vasicola DAR-41379 | 100 | 99.75 | 100 | 99.22 | Xvv | 99.96 | 4.97 | Scaffold | 63.3 | Ensete v GCA_007846285.1 |
| X. vasicola DAR-82733 | 100 | 99.75 | 100 | 99.22 | Xvv | 99.96 | 4.908 | Scaffold | 63.3 | Ensete v GCA_007846135.1 |
| X. vasicola DAR-82572 | 100 | 99.75 | 100 | 99.22 | Xvv | 99.96 | 4.947 | Contig | 63.3 | Ensete v GCA_007846335.1 |
| X. vasicola SY103C | 84 | 67.77 | 94 | 69.24 | BXO8 | 94.3 | 4.662 | Contig | 63.5 | sugarcar GCA_003312635.3 |
| X. vasicola NCPPB 1395 | 100 | 100 | 100 | 100 | Xvv | 99.96 | 4.84 | Scaffold | 63.3 | Ensete v GCA_000772785.1 |
| X. vasicola NCPPB 1241 | 100 | 100 | 100 | 100 | Xvv | 99.96 | 4.851 | Scaffold | 63.3 | Ensete v GCA_000772695.1 |
| X. vasicola NCPPB 1396 | 100 | 100 | 100 | 100 | Xvv | 99.96 | 4.805 | Scaffold | 63.4 | Ensete v GCA_000772725.1 |
| X. vasicola NCPPB 989 | 100 | 99.75 | 100 | 92.6 | Xvv | 99.96 | 4.995 | Scaffold | 63.1 | Ensete v GCA_000772795.1 |
| X. vasicola DAR-82726 | 100 | 100 | 100 | 100 | Xvv | 99.96 | 4.961 | Contig | 63.3 | Ensete v GCA_007846345.1 |
| X. vasicola pv. zeae XZ2 | 100 | 100 | 100 | 99.92 | Xvv | 99.96 | 4.829 | Contig | 63.3 | Ensete v GCA_012844675.1 |
| X. vasicola pv. zeae XZ9 | 100 | 100 | 100 | 99.92 | Xvv | 99.96 | 4.829 | Contig | 63.3 | Ensete v GCA_012844705.1 |
| X. vasicola pv. zeae X09 | 100 | 100 | 100 | 99.92 | Xvv | 99.96 | 4.823 | Contig | 63.3 | Ensete v GCA_012844575.1 |
| X. vasicola pv. zeae X09 | 100 | 100 | 100 | 99.92 | Xvv | 99.96 | 4.486 | Contig | 63.2 | Ensete v GCA_003112065.1 |
| X. vasicola pv. zeae X01 | 100 | 100 | 100 | 99.92 | Xvv | 99.96 | 4.373 | Scaffold | 63.3 | Ensete v GCA_003112105.1 |
| X. vasicola pv. zeae X45 | 100 | 100 | 100 | 99.92 | Xvv | 99.96 | 4.909 | Contig | 63.3 | Ensete v GCA_012844655.1 |
| X. vasicola pv. zeae XGP | 100 | 100 | 100 | 99.92 | Xvv | 99.96 | 4.818 | Contig | 63.3 | Ensete v GCA_012844665.1 |
| X. vasicola pv. zeae X22 | 100 | 100 | 100 | 99.92 | Xvv | 99.96 | 4.96 | Contig | 63.3 | Ensete v GCA_012844585.1 |
| X. vasicola pv. zeae XGP | 100 | 100 | 100 | 99.92 | Xvv | 99.96 | 4.376 | Contig | 63.2 | Ensete v GCA_003111865.1 |
| X. vasicola pv. zeae X15 | 100 | 100 | 100 | 99.92 | Xvv | 99.96 | 4.824 | Contig | 63.3 | Ensete v GCA_012844555.1 |
| X. vasicola pv. zeae X01 | 100 | 100 | 100 | 99.92 | Xvv | 99.96 | 4.944 | Contig | 63.3 | Ensete v GCA_012844535.1 |
| X. vasicola pv. zeae X15 | 100 | 100 | 100 | 99.92 | Xvv | 99.96 | 4.33 | Scaffold | 63.3 | Ensete v GCA_003111965.1 |
| X. vasicola pv. zeae X22 | 100 | 100 | 100 | 99.92 | Xvv | 99.96 | 4.53 | Scaffold | 63.3 | Ensete v GCA_003112025.1 |
| X. vasicola pv. zeae X02 | 100 | 100 | 100 | 99.92 | Xvv | 99.96 | 4.761 | Contig | 63.3 | Ensete v GCA_012844565.1 |
| X. vasicola pv. zeae X02 | 100 | 100 | 100 | 99.92 | Xvv | 99.96 | 4.323 | Scaffold | 63.3 | Ensete v GCA_003112005.1 |
| X. vasicola pv. zeae X23 | 100 | 100 | 100 | 99.92 | Xvv | 99.96 | 4.947 | Contig | 63.3 | Ensete v GCA_012844625.1 |
| X. vasicola pv. zeae X23 | 100 | 100 | 100 | 99.92 | Xvv | 99.96 | 4.439 | Scaffold | 63.3 | Ensete v GCA_003111985.1 |
| X. vasicolaNCPPB 1394 | 100 | 100 | 100 | 100 | Xvv | 99.96 | 4.874 | Scaffold | 63.1 | Ensete v GCA_000774005.1 |
| X. vasicolaDAR-82741 | 100 | 100 | 100 | 100 | Xvv | 99.96 | 4.879 | Scaffold | 63.2 | Ensete v GCA_007846375.1 |
| X. vasicola pv. zeaeXZ2 | 100 | 100 | 100 | 100 | Xvv | 99.96 | 3.849 | Contig | 63.1 | Ensete v GCA_003111845.1 |
| X. vasicola pv. zeaeXZ9 | 100 | 100 | 100 | 100 | Xvv | 99.96 | 3.833 | Contig | 63.2 | Ensete v GCA_003111825.1 |
| X. vasicola pv. zeaeX45 | 100 | 100 | 100 | 100 | Xvv | 99.96 | 3.921 | Contig | 63.2 | Ensete v GCA_003111905.1 |
| X. vasicola NCPPB 902 | 100 | 100 | 100 | 100 | Xvv | 99.96 | 4.916 | Complete | 63.3 | Ensete v GCA_000772775.3 |
| X. vasicola pv. vasculorum SAM11 | 100 | 100 | 100 | 100 | Xvv | 99.96 | 4.871 | Contig | 63.2 | Ensete v GCA_002191955.1 |
| X. vasicola pv. vasculorum SAM-1 | 100 | 100 | 100 | 100 | Xvv | 99.96 | 4.889 | Contig | 63.2 | Ensete v GCA_003725315.1 |
| X. vasicola pv. vasculorum Arg-1A | 100 | 100 | 100 | 100 | Xvv | 99.96 | 4.939 | Contig | 63.2 | Ensete v GCA_002490275.1 |
| X. vasicola pv. vasculorum Arg-4A | 100 | 100 | 100 | 100 | Xvv | 99.96 | 4.947 | Scaffold | 63.2 | Ensete v GCA_003724955.1 |
| X. vasicola pv. vasculorum NCPPB | 100 | 100 | 100 | 100 | Xvv | 99.96 | 5.11 | Contig | 63.3 | Ensete v GCA_019200945.1 |
| X. vasicola pv. vasculorum 645 | 100 | 100 | 100 | 100 | Xvv | 99.96 | 4.882 | Contig | 63.2 | Ensete v GCA_003116615.1 |
| X. vasicola pv. vasculorum 681 | 100 | 100 | 100 | 100 | Xvv | 99.96 | 4.888 | Contig | 63.2 | Ensete v GCA_003116635.1 |
| X. vasicola pv. vasculorum 715 | 100 | 100 | 100 | 100 | Xvv | 99.96 | 4.892 | Contig | 63.2 | Ensete v GCA_003116655.1 |

| | | | | | | | | | | |
|---|---|---|---|---|---|---|---|---|---|---|
| X. vasicola pv. vasculorum NCPPB | 100 | 100 | 100 | 100 | Xvv | 99.96 | 4.901 | Contig | 63.3 | Ensete v GCA_019209805.1 |
| X. vasicola pv. vasculorum NE-448 | 100 | 100 | 100 | 100 | Xvv | 99.96 | 4.924 | Contig | 63.2 | Ensete v GCA_003725255.1 |
| X. vasicola pv. vasculorum KS-444 | 100 | 100 | 100 | 100 | Xvv | 99.96 | 4.918 | Contig | 63.2 | Ensete v GCA_003725135.1 |
| X. vasicola pv. vasculorum NE-413 | 100 | 100 | 100 | 100 | Xvv | 99.96 | 4.947 | Contig | 63.2 | Ensete v GCA_003725915.1 |
| X. vasicola pv. vasculorum NE-8 | 100 | 100 | 100 | 100 | Xvv | 99.96 | 4.934 | Scaffold | 63.2 | Ensete v GCA_003725265.1 |
| X. vasicola pv. vasculorum KS-1 | 100 | 100 | 100 | 100 | Xvv | 99.96 | 4.95 | Scaffold | 63.2 | Ensete v GCA_003725155.1 |
| X. vasicola pv. vasculorum NE-11 | 100 | 100 | 100 | 100 | Xvv | 99.96 | 4.948 | Scaffold | 63.2 | Ensete v GCA_003725875.1 |
| X. vasicola pv. vasculorum IA-2 | 100 | 100 | 100 | 100 | Xvv | 99.96 | 4.931 | Scaffold | 63.2 | Ensete v GCA_003725035.1 |
| X. vasicola pv. vasculorum NE-9 | 100 | 100 | 100 | 100 | Xvv | 99.96 | 4.944 | Scaffold | 63.2 | Ensete v GCA_003725235.1 |
| X. vasicola pv. vasculorum Arg-5B | 100 | 100 | 100 | 100 | Xvv | 99.96 | 4.954 | Scaffold | 63.2 | Ensete v GCA_003724975.1 |
| X. vasicola pv. vasculorum Xvv2 | 100 | 100 | 100 | 100 | Xvv | 99.96 | 4.882 | Contig | 63.2 | Ensete v GCA_010279765.1 |
| X. vasicola pv. vasculorum NE-1 | 100 | 100 | 100 | 100 | Xvv | 99.96 | 4.943 | Scaffold | 63.2 | Ensete v GCA_003724915.1 |
| X. vasicola pv. vasculorum CO-5 | 100 | 100 | 100 | 100 | Xvv | 99.96 | 4.941 | Scaffold | 63.2 | Ensete v GCA_003724995.1 |
| X. vasicola pv. vasculorum CO-6 | 100 | 100 | 100 | 100 | Xvv | 99.96 | 4.926 | Scaffold | 63.2 | Ensete v GCA_003725075.1 |
| X. vasicola pv. vasculorum CO-8 | 100 | 100 | 100 | 100 | Xvv | 99.96 | 4.933 | Scaffold | 63.2 | Ensete v GCA_003725095.1 |
| X. vasicola pv. vasculorum Arg-6B | 100 | 100 | 100 | 100 | Xvv | 99.96 | 4.943 | Scaffold | 63.2 | Ensete v GCA_003725115.1 |
| X. vasicola pv. vasculorum IA-3 | 100 | 100 | 100 | 100 | Xvv | 99.96 | 4.926 | Scaffold | 63.2 | Ensete v GCA_003725165.1 |
| X. vasicola pv. vasculorum NE-4 | 100 | 100 | 100 | 100 | Xvv | 99.96 | 4.936 | Scaffold | 63.2 | Ensete v GCA_003725195.1 |
| X. vasicola pv. vasculorum NE744 | 100 | 100 | 100 | 100 | Xvv | 99.96 | 4.878 | Contig | 63.2 | Ensete v GCA_002191965.1 |
| X. vasicola pv. vasculorum NE-7 | 100 | 100 | 100 | 100 | Xvv | 99.96 | 4.947 | Scaffold | 63.2 | Ensete v GCA_003725215.1 |
| X. vasicola pv. vasculorum Arg-2B | 100 | 100 | 100 | 100 | Xvv | 99.96 | 4.947 | Scaffold | 63.2 | Ensete v GCA_003725055.1 |
| X. vasicola pv. vasculorum SAM-1 | 100 | 100 | 100 | 100 | Xvv | 99.96 | 4.919 | Contig | 63.1 | Ensete v GCA_003725935.1 |
| X. vasicola pv. vasculorum Arg-7A | 100 | 100 | 100 | 100 | Xvv | 99.96 | 4.96 | Scaffold | 63.2 | Ensete v GCA_003725005.1 |
| X. vasicola pv. vasculorum NCPPB | 100 | 100 | 100 | 100 | Xvv | 99.96 | 5.076 | Contig | 63.3 | Ensete v GCA_000278055.2 |
| X. vasicola pv. vasculorum NCPPB | 100 | 100 | 100 | 100 | Xvv | 99.96 | 5.334 | Scaffold | 62.1 | Ensete v GCA_000159795.2 |
| X. vasicola pv. vasculorum NCPPB | 100 | 100 | 100 | 100 | Xvv | 99.96 | 4.804 | Scaffold | 63.2 | Ensete v GCA_000278075.1 |
| X. vasicola pv. vasculorum SAM11 | 100 | 100 | 100 | 100 | Xvv | 99.96 | 4.91 | Complete | 63.2 | Ensete v GCA_003015715.1 |
| X. vasicola pv. vasculorum Xv1601 | 100 | 100 | 100 | 100 | Xvv | 99.96 | 4.957 | Complete | 63.2 | Ensete v GCA_003949975.1 |
| X. vasicola pv. musacearum NCPP | 84 | 67.77 | 94 | 69.24 | BXO8 | 94.3 | 4.731 | Contig | 63.5 | sugarcar GCA_019209735.1 |
| X. vasicola pv. musacearum NCPP | 84 | 67.77 | 94 | 69.24 | BXO8 | 94.3 | 4.746 | Contig | 63.5 | sugarcar GCA_019209705.1 |
| X. vasicola pv. musacearum BCC2 | 100 | 100 | 100 | 100 | Xvv | 99.96 | 4.71 | Scaffold | 63.5 | Ensete v GCA_003932615.1 |
| X. vasicola pv. musacearum E52 | 100 | 100 | 100 | 100 | Xvv | 99.96 | 4.753 | Scaffold | 63.5 | Ensete v GCA_003862455.1 |
| X. vasicola pv. vasculorum NCPPB | 100 | 100 | 100 | 100 | Xvv | 99.96 | 4.826 | Scaffold | 63.3 | Ensete v GCA_000278035.1 |
| X. vasicola pv. vasculorum NCPPB | 100 | 100 | 100 | 100 | Xvv | 99.96 | 4.958 | Scaffold | 63.2 | Ensete v GCA_000278015.1 |
| X. vasicola pv. vasculorum NCPPB | 100 | 100 | 100 | 100 | Xvv | 99.96 | 4.952 | Scaffold | 63.2 | Ensete v GCA_000277995.1 |
| X. campestris pv. musacearum N( | 84 | 67.77 | 94 | 69.24 | BXO8 | 94.3 | 4.705 | Contig | 63.5 | sugarcar GCA_001189905.2 |
| X. campestris pv. musacearum N( | 100 | 100 | 100 | 100 | Xvv | 99.96 | 4.693 | Scaffold | 63.5 | Ensete v GCA_000277875.1 |
| X. campestris pv. musacearum N( | 100 | 100 | 100 | 100 | Xvv | 99.96 | 4.729 | Scaffold | 63.5 | Ensete v GCA_000277955.1 |
| X. campestris pv. musacearum N( | 100 | 100 | 100 | 100 | Xvv | 99.96 | 4.793 | Scaffold | 63.2 | Ensete v GCA_000277975.1 |
| X. campestris pv. musacearum 'K | 100 | 100 | 100 | 100 | Xvv | 99.96 | 4.908 | Scaffold | 63.1 | Ensete v GCA_000233635.1 |
| X. campestris pv. musacearum N( | 100 | 100 | 100 | 100 | Xvv | 99.96 | 4.752 | Scaffold | 63.5 | Ensete v GCA_000277915.1 |
| X. campestris pv. musacearum N( | 100 | 100 | 100 | 100 | Xvv | 99.96 | 4.794 | Scaffold | 61.8 | Ensete v GCA_000159815.1 |
| X. campestris pv. musacearum N( | 100 | 100 | 100 | 100 | Xvv | 99.96 | 4.742 | Scaffold | 63.1 | Ensete v GCA_000277935.1 |
| X. vasicola NCPPB 1060 | 100 | 100 | 100 | 100 | Xvv | 99.96 | 5.064 | Complete | 63.267 | Ensete v GCA_000772715.2 |
| X. campestris pv. musacearum N( | 84 | 67.77 | 94 | 69.24 | BXO8 | 94.3 | 4.838 | Complete | 63.477 | sugarcar GCA_000277895.2 |
| X. vasicola pv. arecae NCPPB 2649 | 100 | 100 | 100 | 100 | Xvv | 99.96 | 4.961 | Complete | 63.387 | Ensete v GCA_000770355.2 |
| X. sacchari CFBP4641 | 100 | 100 | 100 | 100 | Xvv | 78.95 | 4.918 | Contig | 69.1 | Sugarcar GCA_002940085.1 |
| X. sacchari F10 | 100 | 92.33 | 100 | 94 | BXO8 | 81.61 | 4.776 | Scaffold | 69.3 | Sugarcar GCA_014206815.1 |
| X. sacchari LMG 476 | 100 | 100 | 100 | 100 | Xvv | 78.95 | 4.927 | Scaffold | 68.9 | Sugarcar GCA_000831625.1 |
| X. sacchari NCPPB 4393 | 100 | 100 | 100 | 100 | Xvv | 78.95 | 4.898 | Contig | 69 | Sugarcar GCA_000225975.2 |
| X. albilineans CFBP2523 | 100 | 92.68 | 99 | 89.55 | Xvv | 76.38 | 3.684 | Scaffold | 63.1 | Sugarcar GCA_002939705.1 |
| X. albilineans GAB266 | 94 | 75.25 | 98 | 76.54 | Xoc | 75.53 | 3.792 | Scaffold | 62.9 | Sugarcar GCA_000963065.1 |
| X. albilineans FIJ080 | 100 | 92.68 | 99 | 89.55 | Xvv | 76.38 | 3.681 | Scaffold | 63 | Sugarcar GCA_000962995.1 |
| X. albilineans MTQ032 | 100 | 92.68 | 100 | 89.55 | Xvv | 76.38 | 3.802 | Scaffold | 62.9 | Sugarcar GCA_000963075.1 |
| X. albilineans HVO082 | 94 | 75.25 | 98 | 76.54 | Xoc | 75.53 | 3.636 | Scaffold | 63 | Sugarcar GCA_000962915.1 |
| X. albilineans REU209 | 94 | 75.25 | 98 | 76.54 | Xoc | 75.53 | 3.687 | Scaffold | 63 | Sugarcar GCA_000963135.1 |
| X. albilineans GPE PC86 | 100 | 92.25 | 100 | 89.55 | Xvv | 76.38 | 3.851 | Scaffold | 62.8 | Sugarcar GCA_000963025.1 |
| X. albilineans XaFL07-1 | 100 | 92.25 | 100 | 89.55 | Xvv | 76.38 | 3.799 | Scaffold | 62.9 | Sugarcar GCA_000963195.1 |
| X. albilineans GPE PC17 | 100 | 92.25 | 100 | 89.55 | Xvv | 76.38 | 3.811 | Scaffold | 62.8 | Sugarcar GCA_000962935.1 |
| X. albilineans REU174 | 100 | 92.25 | 100 | 89.55 | Xvv | 76.38 | 3.863 | Scaffold | 62.8 | Sugarcar GCA_000963115.1 |
| X. albilineans HVO005 | 94 | 75.25 | 98 | 76.54 | Xoc | 75.53 | 3.653 | Scaffold | 62.9 | Sugarcar GCA_000963055.1 |
| X. albilineans USA048 | 100 | 92.25 | 100 | 89.55 | Xvv | 76.38 | 3.582 | Scaffold | 63.1 | Sugarcar GCA_000963145.1 |
| X. albilineans Xa23R1 | 100 | 92.25 | 100 | 89.55 | Xvv | 76.38 | 3.549 | Scaffold | 63.1 | Sugarcar GCA_000963155.1 |

| | | | | | | | | | | |
|---|---|---|---|---|---|---|---|---|---|---|
| X. albilineans LKA070 | 100 | 92.25 | 99 | 89.55 | Xvv | 76.38 | 3.668 | Scaffold | 63.1 | Sugarcar | GCA_000962945.1 |
| X. albilineans PNG130 | 94 | 75.25 | 98 | 76.54 | Xoc | 75.53 | 3.543 | Scaffold | 63.3 | Sugarcar | GCA_000962925.1 |
| X. albilineans Xa-FJ1 | 100 | 92.25 | 100 | 89.55 | Xvv | 76.38 | 3.756 | Complete | 62.974 | Sugarcar | GCA_009931595.1 |
| X. albilineans GPE PC73 | 100 | 92.25 | 100 | 89.55 | Xvv | 76.38 | 3.852 | Complete | 62.906 | Sugarcar | GCA_000087965.1 |
| Xanthomonas citri pv. citri strain | 100 | 96.63 | 100 | 100 | Xoc | 100 | 5.386 | Complete | 64.598 | citrus pla | GCA_016801615.1 |
| Xanthomonas citri pv. citri strain | 100 | 96.63 | 100 | 100 | Xoc | 100 | 5.552 | Complete | 64.738 | citrus pla | GCA_016801695.1 |
| Xanthomonas citri pv. citri strain | 100 | 96.63 | 100 | 100 | Xoc | 100 | 5.596 | Complete | 64.639 | citrus pla | GCA_016801655.1 |
| Xanthomonas citri pv. citri strain | 100 | 96.63 | 100 | 100 | Xoc | 100 | 5.371 | Complete | 64.598 | citrus pla | GCA_016801635.1 |
| Xanthomonas citri pv. citri strain | 100 | 96.63 | 100 | 100 | Xoc | 100 | 5.551 | Complete | 64.733 | citrus pla | GCA_016801675.1 |
| Xanthomonas citri pv. citri strain | 100 | 96.63 | 100 | 100 | Xoc | 100 | 5.51 | Complete | 64.741 | citrus pla | GCA_016801715.1 |
| Xanthomonas citri pv. citri strain | 100 | 96.63 | 100 | 100 | Xoc | 100 | 5.575 | Complete | 64.618 | citrus pla | GCA_002139955.1 |
| Xanthomonas citri pv. citri strain | 100 | 96.63 | 100 | 100 | Xoc | 100 | 5.398 | Complete | 64.664 | citrus pla | GCA_000961435.1 |
| Xanthomonas citri pv. citri strain | 100 | 96.63 | 100 | 100 | Xoc | 100 | 5.398 | Complete | 64.664 | citrus pla | GCA_000961495.1 |
| Xanthomonas citri pv. citri strain | 100 | 96.63 | 100 | 100 | Xoc | 100 | 5.398 | Complete | 64.664 | citrus pla | GCA_000961475.1 |
| Xanthomonas citri pv. citri strain | 100 | 96.63 | 100 | 100 | Xoc | 100 | 5.398 | Complete | 64.664 | citrus pla | GCA_000961455.1 |
| Xanthomonas citri subsp. citri Aw | 100 | 96.63 | 100 | 100 | Xoc | 100 | 5.399 | Complete | 64.664 | citrus pla | GCA_000349225.1 |
| Xanthomonas citri pv. citri strain | 100 | 96.63 | 100 | 100 | Xoc | 100 | 5.514 | Complete | 64.646 | citrus pla | GCA_002139995.1 |
| Xanthomonas citri pv. citri strain | 100 | 96.63 | 100 | 100 | Xoc | 100 | 5.501 | Complete | 64.649 | citrus pla | GCA_002139975.1 |
| Xanthomonas fragariae strain YL1 | 100 | 68.85 | 100 | 100 | Xoc | 94 | 4.11 | Complete | 61.99 | strawber | GCA_017603965.1 |
| Xanthomonas sp. SI chromosome | 100 | 68.85 | 100 | 100 | Xoc | 77 | 5.25 | Complete | 68.4 | Ryegrass | GCA_014236855.1 |
| Xanthomonas sp. SS chromosome | 100 | 68.85 | 100 | 100 | Xoc | 77 | 5.19 | Complete | 68.5 | Ryegrass | GCA_014236835.1 |
| Xanthomonas arboricola pv. prun | 100 | 68.85 | 100 | 100 | Xoc | 54 | 5.14 | Complete | 65.4 | Hazelnut | GCA_008761555.1 |
| Xanthomonas arboricola pv. cory | 100 | 68.85 | 100 | 100 | Xoc | 54 | 5.23 | Complete | 65.38 | Hazelnut | GCA_905220805.1 |
| Xanthomonas arboricola pv. cory | 100 | 68.85 | 100 | 100 | Xoc | 54 | 5.08 | Complete | 65.6 | Hazelnut | GCA_905220785.1 |
| Xanthomonas arboricola pv. cory | 100 | 68.85 | 100 | 100 | Xoc | 54 | 5.29 | Complete | 65.38 | Hazelnut | GCA_905220715.1 |
| Xanthomonas arboricola pv. cory | 100 | 68.85 | 100 | 100 | Xoc | 54 | 5.17 | Complete | 65.5 | Hazelnut | GCA_018141705.1 |
