## supplementary figure 1, 2, 3 for "Systematic hyper-variation and evolution at a lipopolysaccharide locus in the population of *Xanthomonas* species that infect rice and sugarcane"

**Supplementary figures**

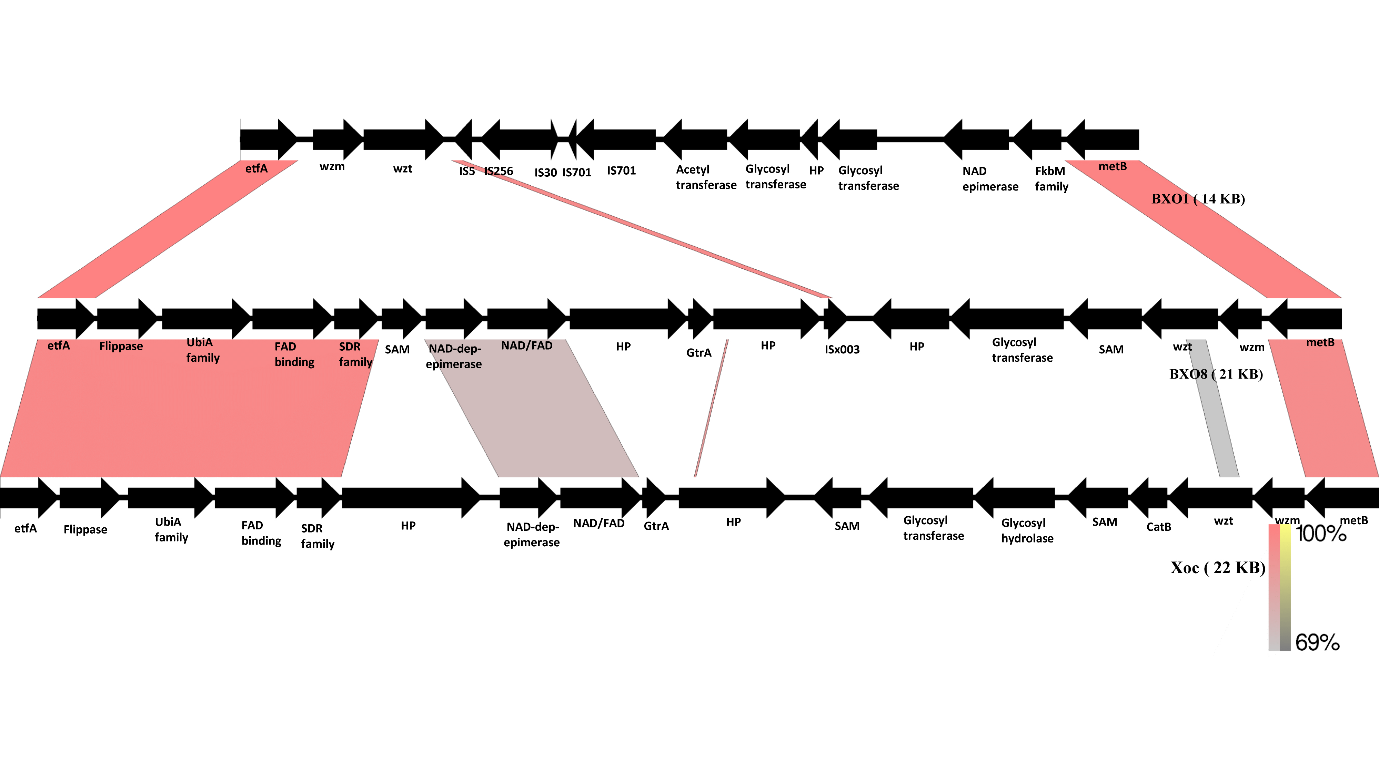
**Supplementary figure 1:** LPS synteny comparison of BXO1, BXO8 and Xoc type LPS cassette with Easyfig and the BLASTn algorithm. Arrows represent the location of genes and shaded lines reflect the degree of homology between pairs of genes in two LPS cassettes.

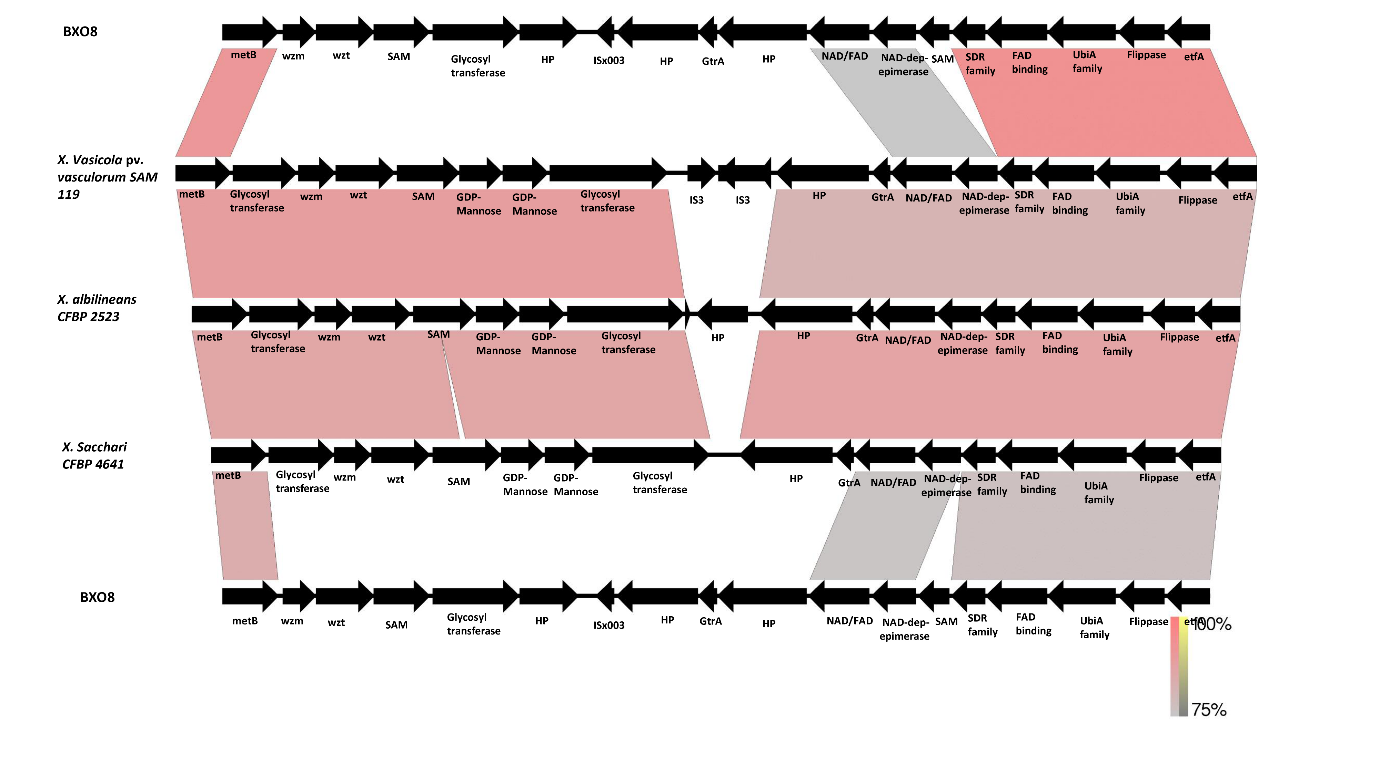
**Supplementary figure 2:** LPS synteny comparison with Easyfig and the BLASTn algorithm. Arrows represent the location of genes and shaded lines reflect the degree of homology between pairs of genes in two LPS cassettes. *X. sacchari* CFBP 4641 and *X. albilineans* CFBP 2523 shows more homology with Xvv type of LPS than BXO8.

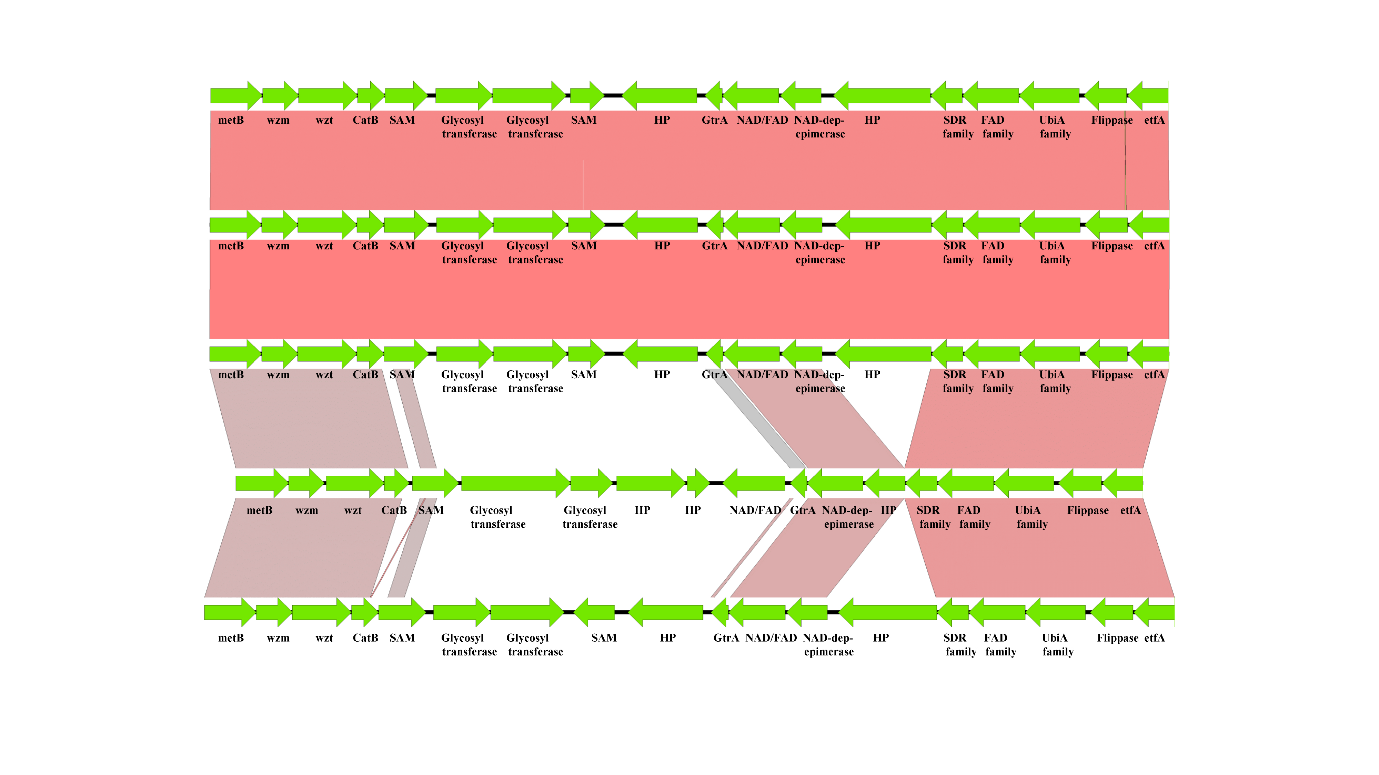
**Supplementary figure 3:** LPS synteny comparison with Easyfig and the BLASTn algorithm. Arrows represent the location of genes and shaded lines reflects the degree of homology between pairs of genes in two LPS cassettes. Gene cluster in Xcc strains (DAR84832, AW13), *X. fragariae* YL19, *X. arboricola* pv. *corylina* A7 LPS showing homology with Xoc type LPS cassette (green color).
